# Confidence as a noisy decision reliability estimate

**DOI:** 10.1101/2021.12.17.473249

**Authors:** Zoe M. Boundy-Singer, Corey M. Ziemba, Robbe L. T. Goris

## Abstract

Decisions vary in difficulty. Humans know this and typically report more confidence in easy than in difficult decisions. However, confidence reports do not perfectly track decision accuracy, but also reflect response biases and difficulty misjudgments. To isolate the quality of confidence reports, we developed a model of the decision-making process underlying choice-confidence data. In this model, confidence reflects a subject’s estimate of the reliability of their decision. The quality of this estimate is limited by the subject’s uncertainty about the uncertainty of the variable that informs their decision (“meta-uncertainty”). This model provides an accurate account of choice-confidence data across a broad range of perceptual and cognitive tasks, revealing that meta-uncertainty varies across subjects, is stable over time, generalizes across some domains, and can be manipulated experimentally. The model offers a parsimonious explanation for the computational processes that underlie and constrain the sense of confidence.

---

Humans are aware of the fallibility of perception and cognition. When we experience a high degree of confidence in a perceptual or cognitive decision, that decision is more likely to be correct than when we feel less confident ^1^. This “metacognitive” ability helps us to learn from mistakes ^2^, to plan future actions ^3^, and to optimize group decision-making ^4^. There is a long-standing interest in the mental operations underlying our sense of confidence ^5–7^, and the rapidly expanding field of metacognition seeks to understand how metacognitive ability varies across domains ^8^, individuals ^9^, clinical states ^10^, and development ^11^.

Quantifying a subject’s ability to introspect about the correctness of a decision is a challenging problem ^12–14^. There exists no generally agreed-upon method ^15^. Even in the simplest decision-making tasks, several distinct factors influence a subject’s confidence reports. Consider a subject jointly reporting a binary decision about a sensory stimulus (belongs to “Category A” vs “Category B”) and their confidence in this decision. Confidence reports will reflect the subject’s ability to discriminate between both stimulus categories – the higher this ability, the higher the reported confidence ^16^. They will also reflect the subject’s response bias (e.g., a large willingness to declare “high confidence” or “Category A”) ^17–19^. Yet, neither of these factors characterizes the subject’s metacognitive ability ^13^.

Here, we introduce a method to quantify metacognitive ability on the basis of choice-confidence data. We propose that confidence reflects a subject’s estimate of the reliability of their decision ^20^, expressed in units of signal-to-noise ratio. This estimate results from a computation involving the uncertainty of the decision variable that informed the subject’s choice ^21^. It follows that metacognitive ability is determined by the subject’s knowledge about this uncertainty, or lack thereof (i.e., uncertainty about uncertainty, hereafter termed “meta-uncertainty”). The more certain a subject is about the uncertainty of the decision variable, the lower their meta-uncertainty, and the better they are able to assess the reliability of a decision. We leverage modern computational techniques to formalize this hypothesis in a two-stage process model that is rooted in traditional signal detection theory ^22^ and that can be fit to choice-confidence data (the “CASANDRE” or “Confidence AS A Noisy Decision Reliability Estimate” model). The model predicts a systematic dependency of confidence on choice consistency ^20^ and naturally separates metacognitive ability from discrimination ability and response bias.

We found that this model provides an excellent account of choice-confidence data reported in a large set of previously published studies ^23–28^. Our analysis suggests that meta-uncertainty provides a better metric for metacognitive ability than the non-process-model based alternatives that currently prevail in the literature ^13,15^. Specifically, meta-uncertainty has higher test-retest reliability, is less affected by discrimination ability and response bias, and has comparable cross-domain generalizability. Meta-uncertainty is higher in tasks that involve more levels of stimulus uncertainty, implying that it can be manipulated experimentally. Together, these results illuminate the mental operations that give rise to our sense of confidence, and they provide evidence that metacognitive ability is fundamentally limited by subjects’ uncertainty about the reliability of their decisions.

## Results

In simple decision-making tasks, human confidence reports lawfully reflect choice consistency ^20^. Consider two example subjects who performed a two-alternative forced choice (2-AFC) categorization task in which they judged on every trial whether a visual stimulus belonged to category A or B, and additionally reported their confidence in this decision using a four-point rating scale. Categories were characterized by distributions of stimulus orientation that were predominantly smaller (A) or larger (B) than zero degrees. Stimuli varied in orientation and contrast (Fig. 1a). Because the category distributions overlap, errors are inevitable. The most accurate strategy is to choose category A for all stimuli whose orientation is smaller than zero degrees, and category B for all stimuli whose orientation exceeds zero degrees (Fig. 1b, top, dotted line). As can be seen from the aggregated choice behavior, the more the stimulus orientation deviates from zero, the more closely human subjects approximate this ideal (Fig. 1b, top, symbols). As can also be seen, this relationship is modulated by stimulus contrast – the lower the stimulus contrast, the weaker the association between orientation and choice (Fig. 1b, top, green vs yellow symbols). The distinct effects of orientation and contrast on choice consistency are evident in the subjects’ confidence reports. Confidence is minimal for conditions associated with a choice proportion near 0.5 (i.e., the most difficult conditions), and monotonically increases as choice proportions deviate more from 0.5 (Fig. 1b, bottom). We found that the association between choice consistency and confidence is so strong, that plotting average confidence level against the aggregated choice behavior reveals a single relationship across all stimulus conditions (Fig. 1c). This is true of both example subjects, although their confidence-consistency relationships differ in shape, offset, and range. We speculate that a lawful confidence-consistency relationship is not a coincidental feature of this experiment, but a widespread phenomenon in confidence studies (Fig. 1d).

**Figure 1.**
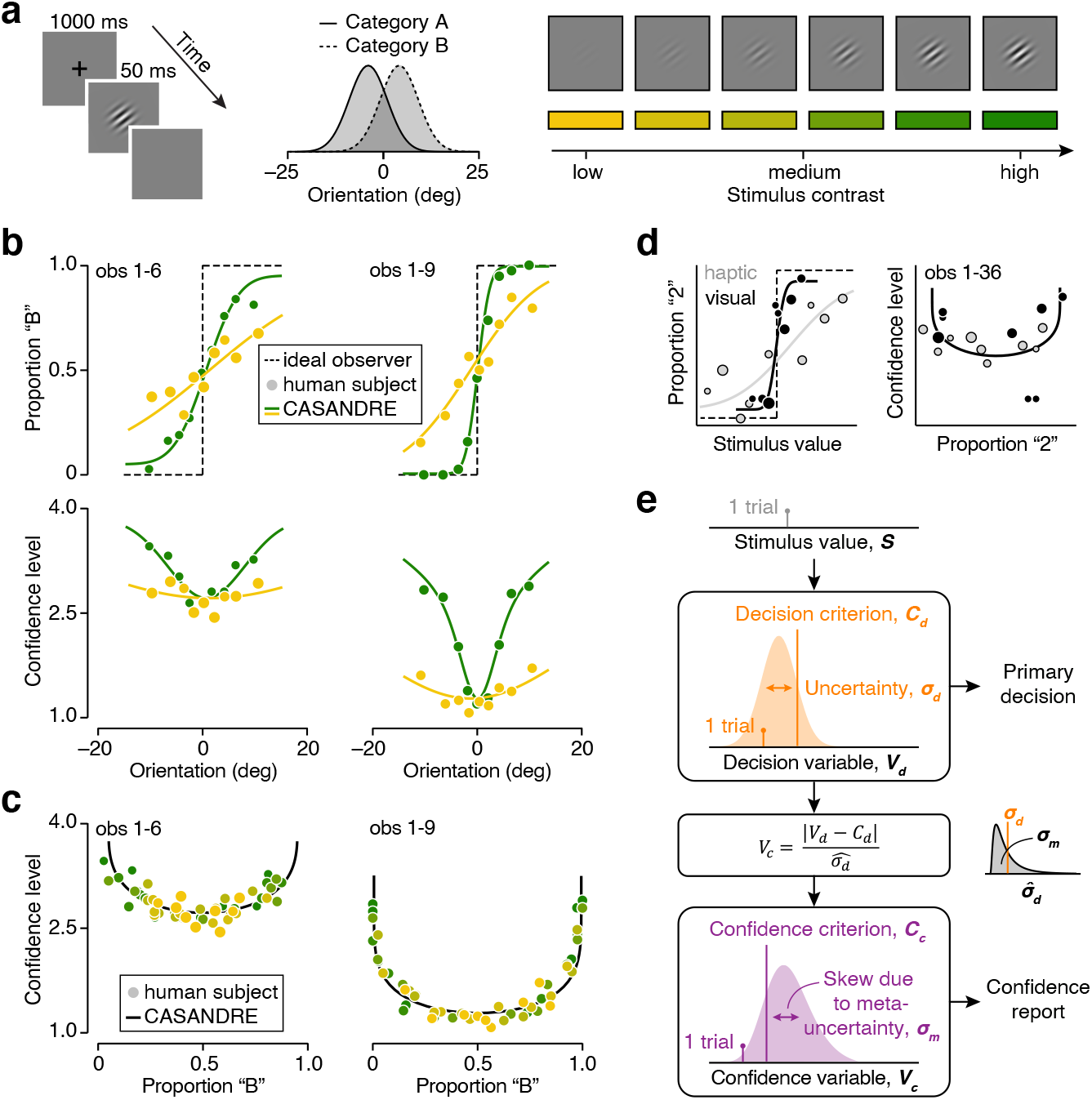
CASANDRE, a two-stage process model of decision confidence, accounts for the relation between confidence reports and choice consistency. (**a**) Experimental design employed by ref. ^25^. (**b**) Top: Proportion of “Category B” choices is plotted against stimulus orientation, split out by stimulus contrast (green vs yellow), for two example subjects (left, obs 1-6: Observer 6 in experiment 1 from ref. ^25^; right, obs 1-9: Observer 9 in experiment 1 from ref. ^25^). Bottom: Same for mean confidence level. Symbols summarize observed choice behavior, the dotted line illustrates the theoretical optimum, and the full lines show the fit of the CASANDRE model. Symbol size is proportional to the number of trials. The model was fit to all data simultaneously using a maximum likelihood estimation method. Only two out of six contrasts are shown here. Fits to all conditions are shown in Supplementary Fig. 1. (**c**) Observed and predicted confidence-consistency relationship for two example subjects. (**d**) Observed and predicted choice-confidence data for an example subject performing a visuo-haptic two-interval forced choice (2-IFC) categorization task (observer 36 in experiment 1 from Arbuzova and Filevich in the Confidence Database ^23^). (**e**) Schematic of the hierarchical decision-making process underlying choice-confidence data in the CASANDRE model.

A monotonically increasing relation between confidence reports and choice consistency implies that subjects can assess the reliability of their decisions. However, whether their ability to do so is excellent or poor cannot be deduced from empirical measurements alone. One possibility is that subjects accurately assess decision reliability on every single trial, indicating excellent metacognitive ability. Alternatively, there might be a high degree of cross-trial variability in confidence reports, implying less accurate decision reliability assessment and thus limited metacognitive ability. Of course, given the variability of the primary choice behavior, some variability in confidence reports is expected, even for flawless introspection. How much exactly? And what might be the origin of excess variance? Answering these questions requires a quantitative model that provides an analogy for the mental operations that underlie a subject’s primary decisions and confidence reports. In the following section, we develop such a process model.

### A two-stage process model of decision-making

Assume that a subject solves a binary decision-making task by comparing a noisy, one-dimensional decision variable, *V*_*d*_, to a fixed criterion, *C*_*d*_ (Fig. 1e, top). For some tasks, it is convenient to think of this decision variable as representing a direct estimate of a stimulus feature (e.g., orientation for the task shown in Fig. 1a). For other tasks, it is more appropriate to think of it as representing the accumulated evidence that favors one response alternative over the other (e.g., “Have I heard this song before?”). The process model specified by these assumptions has proven very useful in the study of perception and cognition. It readily explains why repeated presentations of the same stimulus often elicit variable choices. In doing so, it clarifies how choices reflect a subject’s underlying ability to solve the task as well as their primary response bias ^22^.

We expand this framework with an analogous second processing stage that informs the subject’s confidence report. Assume that the subject is presented with a set of stimuli that elicit the same level of cross-trial variability in the decision variable. The smaller the overlap of the stimulus-specific decision variable distribution with the decision criterion, the “stronger” the associated stimulus is, and the more consistent choices will be. On any given trial, the distance between the decision variable and the decision criterion provides an instantaneously available proxy for stimulus strength, and hence for choice reliability ^14,29–33^. However, in many tasks, the decision variable’s dispersion, *σ*_*d*_, will vary across conditions, resulting in different amounts of stimulus “uncertainty” (the larger *σ*_*d*_, the greater this uncertainty). To be a useful proxy for choice reliability, the stimulus strength estimate must therefore be normalized by this factor ^20^. This operation yields a unitless, positive-valued variable, *V*_*c*_, which represents the subject’s confidence in the decision:

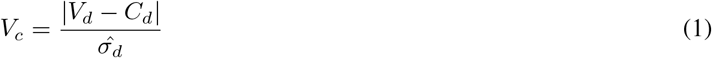

where *V*_*d*_ is the decision variable, *C*_*d*_ the decision criterion, and 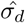 the subject’s estimate of *σ*_*d*_. We assume that the subject is unsure about the exact level of stimulus uncertainty. Repeated trials will thus not only elicit different values of the decision variable, but will also elicit different estimates of stimulus uncertainty. Specifically, we assume that 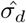 is on average correct (i.e., its mean value equals *σ*_*d*_), but varies from trial-to-trial with standard deviation *σ*_*m*_, resulting in “meta-uncertainty” (the larger *σ*_*m*_, the greater this meta-uncertainty). As we shall see, variability in the decision variable is the critical model component that limits stimulus discriminability, while variability in the uncertainty estimate similarly limits metacognitive ability. Finally, comparing the confidence variable with a fixed criterion, *C*_*c*_, yields a confidence report (Fig. 1e, bottom).

To fit this model to data, the form of the noise distributions must be specified. A common choice for the first-stage noise is the normal distribution. This choice is principled, as the normal distribution is the maximum entropy distribution for real-valued signals with a specified mean and variance ^34^. It is also convenient, as it results in fairly simple data-analysis recipes ^22^. The second-stage noise describes variability of a positive-valued signal (*σ*_*d*_ cannot be smaller than zero by definition). A suitable maximum entropy distribution for such a variable is the log-normal distribution ^28,34^. Under these assumptions, the confidence variable is a probability distribution constructed as the distribution of the ratio of a normally and log-normally distributed variable. There exists no closed form description of this ratio distribution, ruling out simple data-analysis recipes. However, we can leverage modern computational tools to quickly compute the confidence variable’s probability density function by describing it as a mixture of Gaussian distributions (see Methods). This mathematical street-fighting maneuver ^35^ enables us to fit this two-stage process model to choice data (Fig. 1b-d, full lines). Before doing so, we first derive a set of basic model predictions.

### Deriving model predictions

To gain a deeper understanding of the impact of the different model components on confidence reports, we investigated the model’s behavior in a continuous 2-AFC discrimination task with binary confidence report options (“confident” or “not confident”). We assumed the decision variable’s mean value to be stimulus-dependent (in this simulation, it is identical to the true stimulus value). All other model components were varied independently of the stimulus (see Methods). Altering the first-stage decision criterion (Fig. 2a, top left, orange vs grey line) affects the confidence variable distribution by shifting its mode and, in the presence of meta-uncertainty, its spread and skew (Fig. 2a, bottom left, purple vs grey distribution). At the level of observables, this manipulation results in a horizontal shift of the “psychometric function” that characterizes how choices depend on stimulus value (Fig. 2a, top right). This shift is accompanied by an identical shift of the “confidence function” that characterizes how confidence reports depend on stimulus value (Fig. 2a, bottom right). Effects of this kind have been documented for human ^20,36,37^ and animal ^38,39^ subjects. Altering the level of first-stage noise (Fig. 2b, top left, orange vs grey distribution) affects the confidence variable distribution by changing its mode and, in the presence of meta-uncertainty, its spread and skew (Fig. 2b, bottom left, purple vs grey distribution). At the level of choice behavior, this manipulation changes the slope of the psychometric function (Fig. 2b, top right) as well as the overall fraction of “confident” reports (Fig. 2b, bottom right). In contrast, the parameters that control the model’s second-stage operations do not affect the primary choice behavior but only the confidence reports. Specifically, changing the confidence criterion (Fig. 2c, bottom left, purple vs grey lines) mainly impacts the confidence function by shifting it vertically (Fig. 2c, bottom right). Changing the level of meta-uncertainty alters the confidence variable distribution’s mode, variance, and skew (Fig. 2d, bottom left, purple vs grey distribution), resulting in a complex pattern of changes in the confidence function (Fig. 2d, bottom right).

**Figure 2.**
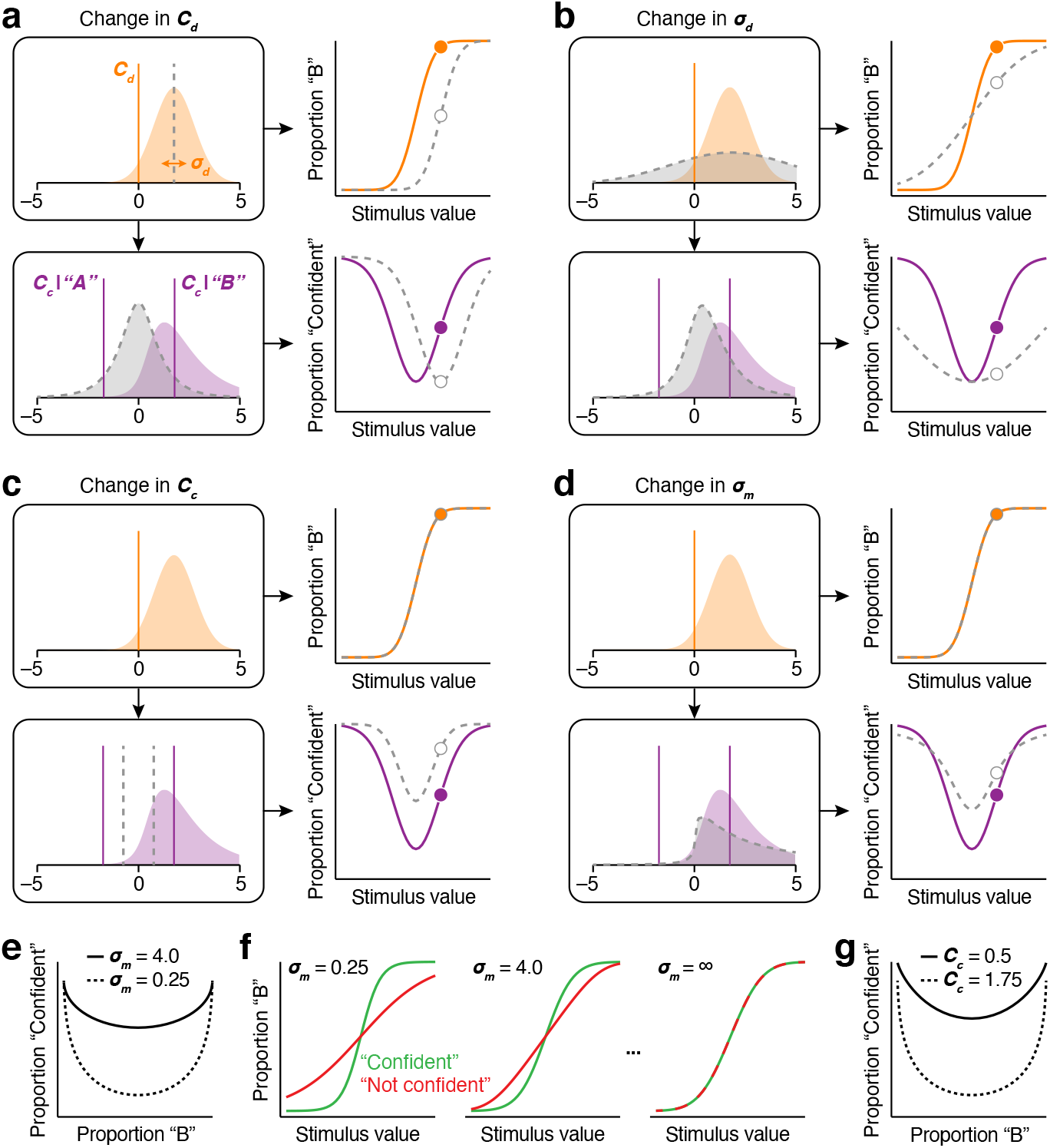
Impact of the different model components on primary choice behavior and confidence reports. (**a**) Top left: illustration of the decision criterion (orange line) and the decision variable distribution elicited by repeated presentations of the same stimulus (orange distribution). Bottom left: the associated confidence variable distribution (purple distribution). *V*_*c*_ is a positive-valued variable. As plotting convention, we reserve negative values for “Category A” choices, and positive values for “Category B” choices. The confidence criterion (purple line) therefore shows up twice in this graph. Top right: the resulting psychometric function over a range of stimulus values (orange line). The filled symbol corresponds to the condition depicted on the left hand side. Bottom right: same for the resulting confidence function. All panels: the grey dotted line illustrates how the model predictions change when a specific model component (here, the decision criterion) is altered. The open symbol corresponds to the condition depicted on the left hand side. (**b**) Increasing the level of stimulus uncertainty affects both primary decisions and confidence reports. (**c**) Lowering the confidence criterion yields more “confident” reports at all stimulus values. (**d**) Increasing meta-uncertainty increases the fraction of “confident” reports for weak stimuli, but has the opposite effect for strong stimuli. (**e**) The confidence-consistency relation for two levels of meta-uncertainty. All other model parameters held equal. (**f**) The psychometric function, split out by confidence report (“confident” in green vs “not confident” in red), for three levels of meta-uncertainty. (**g**) The confidence-consistency relation under a liberal vs a conservative confidence criterion. All other model parameters held equal, *σ*_*m*_ = 0.25.

What does it mean to say that someone has good or bad self-knowledge? The CASANDRE model provides a principled answer that is at once intuitive and revealing. Everything held equal, increasing meta-uncertainty makes the confidence variable distribution more heavy-tailed (Fig. 2d, bottom left). This in turn leads to an increase in the fraction of “confident” reports for weak stimuli, but has the opposite effect for strong stimuli (Fig. 2d, bottom right). As a consequence, the dynamic range of the confidence-consistency relation decreases (Fig. 2e). However, these effects are not balanced. In particular, when meta-uncertainty is high, there is a dramatic increase in “confident” reports for the most difficult conditions (Fig. 2e, full black line). This increase does not reflect an actual change in task performance (Fig. 2d, top right). Rather, the association between confidence and choice consistency has weakened. This can be appreciated by inspecting the psychometric function split out by confidence report. When meta-uncertainty is low, “confident” decisions tend to be much more reliable than “not confident” decisions (Fig. 2f, left, green vs red). As meta-uncertainty increases, this distinction weakens and eventually disappears (Fig. 2f, middle-right). In sum, under the CASANDRE model, a lack of self-knowledge means having a limited capacity to distinguish reliable from unreliable decisions (note that this is not the same as distinguishing correct from incorrect decisions) ^20^. This is a practical and useful insight. However, the magnitude of the effects shown in Fig. 2e,f depends on the other model components as well (e.g., Fig. 2g). These components will rarely be constant across tasks, individuals, or the life-span. Determining the level of meta-uncertainty therefore requires directly fitting the model to data.

### Evaluating the model architecture

We have motivated our framework on the basis of a qualitative observation (the lawful confidence-consistency relationship) and first principles (the inherent noisiness of perceptual and cognitive processes). To further test the central tenets of the CASANDRE model, we quantitatively examined the choice-confidence data collected by Adler and Ma (2018). We conducted several model comparisons designed to interrogate the framework’s second-stage operations. For this reason, we began by fitting the first-stage parameters to each subject’s choice data and then kept these parameters constant across all model variants (see example in Fig. 3a). We first asked whether a simpler computation can account for confidence reports. We compared a model variant in which confidence reflects a subject’s estimate of stimulus strength ^14,29–33^ with one in which it reflects an estimate of decision reliability (i.e., stimulus strength normalized by stimulus uncertainty; Fig. 3b, left). To quantify model quality, we computed each model’s AIC value (see Methods). For all 19 subjects, the more complex model outperformed the simpler variant (median difference in AIC = 1179.5; Fig. 3c, top). We then asked whether meta-uncertainty is a necessary model component, and found this to be the case (Fig. 3b, middle). Including meta-uncertainty improved model quality for all 19 subjects (median difference in AIC = 285.2; Fig. 3c, middle). These model comparisons thus provide strong and consistent support for the hypothesis that confidence reflects a subject’s noisy estimate of the reliability of their decision.

**Figure 3.**
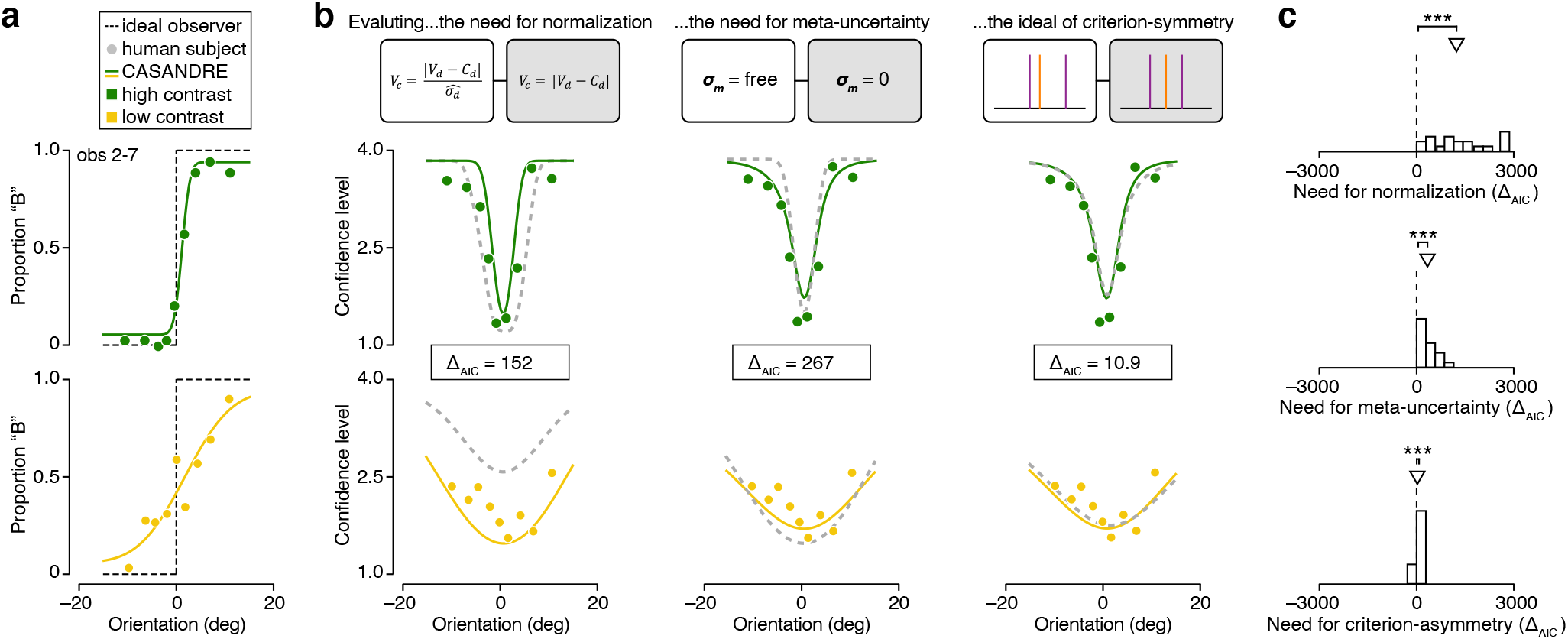
Comparison of different model architectures. (**a**) Proportion of “Category B” choices is plotted against stimulus orientation for high and low contrast stimuli (top vs bottom). Symbols summarize observed choice behavior of an example subject (observer 7 in experiment 2 from ref. ^25^), the dotted line illustrates the theoretical optimum, and the full lines show the fit of the first stage which is shared across all model variants examined in this analysis. As previously, the model was fit to all data simultaneously. (**b**) Mean confidence level is plotted as a function of stimulus orientation for the example subject. (Left) Fits of two model variants in which confidence either reflects an estimate of decision reliability (full lines) or of stimulus strength (dashed lines). (Middle) Fits of two model variants in which confidence either reflects a noisy (full lines) or noiseless (dashed lines) estimate of decision reliability. (Right) Fits of two model variants in which confidence criteria either depend (full lines) or not (dashed lines) on the primary decision. (**c**) Distribution of the difference in AIC value for each model comparison across 19 subjects. Positive values indicate evidence for the more complex model variant. Arrows indicate median of the distribution. *** P < 0.001, Wilcoxon signed-rank test.

Further attempts to improve the model architecture yielded comparatively weak and inconsistent results. In particular, we wondered whether model performance would benefit from allowing criterion-asymmetry (meaning that the confidence criteria depend on the primary decision) and adopting a different second-stage noise distribution (the Gamma distribution). Allowing criterion-asymmetry improved model performance for 16 out of 19 subjects (median difference in AIC = 27.9; Fig. 3b, right; Fig. 3c, bottom; different example subject shown in Supplementary Fig. 2), while the log-normal distribution was preferred over the Gamma distribution for 16 out of 19 subjects (median difference in AIC = 17.7). For simplicity, we chose to use symmetric confidence criteria for all further analyses. Finally, we compared the CASANDRE model with a model recently proposed by Shekhar and Rahnev (2021; the “criteria-noise model”). In this model, confidence reflects a subject’s estimate of evidence strength and metacognitive ability is limited by a subject’s inability to maintain constant confidence criteria across trials ^28^. As this model is tailored to experiments that employ only two levels of stimulus strength, we examined the choice-confidence data collected by Shekhar and Rahnev (2021) and found that the CASANDRE model either matched or outperformed the criteria-noise model (Supplementary Fig. 8a; see Supplementary Information).

### Estimating meta-uncertainty from sparse data

We seek to quantify a subject’s ability to introspect about the reliability of a decision. Our method consists of interpreting human choice-confidence data through the lens of a principled two-stage process model. What kind of measurements are required to obtain robust and reliable estimates of meta-uncertainty, the model’s parameter that governs metacognitive ability? We verified that Adler and Ma’s experimental design affords solid parameter recovery (See Supplementary Fig. 3). However, their design is exceptional for its large number of stimulus conditions ^25^. Many studies use as little as two conditions ^23^. To test whether our approach generalizes to such experiments, we performed a recovery analysis. We used the CASANDRE model to generate synthetic data sets for five model subjects performing a 2-AFC discrimination task with binary confidence report options (see Methods). The model subjects only differed in their level of meta-uncertainty, which ranged from negligible to considerable (Fig. 4a, colored lines). We simulated data for each model subject using experimental designs that varied in the number of trials (100 vs 1,000) and in the number of conditions (2 vs 20; Fig. 4a, top). Figure 4b summarizes an example synthetic experiment. The model parameters (*σ*_*d*_, *C*_*d*_, *σ*_*m*_, *C*_*c*_) specify the relation between stimulus value and the probability of each response option (Fig. 4b, left). We used these probabilities to simulate a synthetic dataset of 1,000 trials distributed across 20 conditions (Fig. 4b, middle). We then identified the set of parameter values that best describes these data (Fig. 4b, right). We repeated this procedure 100 times for each simulated experiment. Our method yields robust estimates of meta-uncertainty: for all model subjects and all experimental designs, the median estimate closely approximates the ground truth value (Fig. 4c, symbols). The reliability of these estimates is higher for more trials and somewhat higher for denser stimulus sampling (Fig. 4c, error bars). Estimation error in *σ*_*m*_ covaried with estimation error in *C*_*c*_ (Supplementary Fig. 7). We conclude that the CASANDRE model typically can be identified in sparse experimental designs.

**Figure 4.**
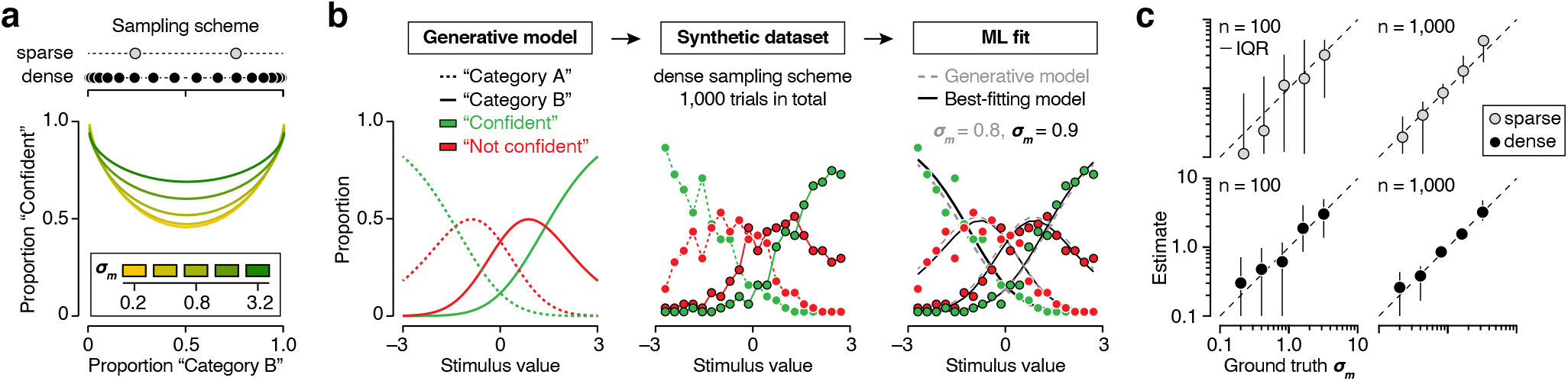
Model recovery analysis. (**a**) We simulated choice-confidence data for five model subjects who differed in their level of meta-uncertainty (colored lines) using experimental designs that varied in the number of trials (100 vs 1,000) and in the number of conditions (2 vs 20, grey and black symbols). (**b**) An example synthetic experiment and model-based analysis. (**c**) The median estimate of meta-uncertainty is plotted against the ground truth value for four experimental designs. Meta-uncertainty was limited to a minimum value of 0.1. Error bars illustrate the interquartile range (IQR) computed from 100 simulated data sets.

### Meta-uncertainty: construct reliability and validity

So far, we have presented evidence that confidence is well described as reflecting a subject’s decision-reliability estimate. In the CASANDRE model, the quality of this estimate is limited by meta-uncertainty. This naturally raises the question of whether meta-uncertainty is a “real” thing. In other words, do meta-uncertainty estimates isolate a stable property of human subjects that captures their metacognitive ability?

The most straightforward form of stability is repeatability. If we were to measure a subject’s meta-uncertainty on two different occasions using the same experimental paradigm, we should obtain similar estimates. Navajas et al. (2017) conducted a perceptual confidence experiment in which 14 subjects performed the same task twice with approximately one month in between test sessions ^24^. We used the CASANDRE model to analyze their data (see Methods and Supplementary Fig. 4). Measured and predicted choice-confidence data were strongly correlated, indicating that the model describes the data well (condition-specific proportion correct choices: Spearman’s rank correlation coefficient *r* = 0.96, *P* < 0.001; condition-specific mean confidence level: *r* = 0.99, *P* < 0.001). Critically, we found meta-uncertainty estimates to be strongly correlated across both sessions as well (*r* = 0.78, *P* = 0.002; Fig. 5a). This suggests that meta-uncertainty measures a stable characteristic of human confidence reporting behavior.

**Figure 5.**
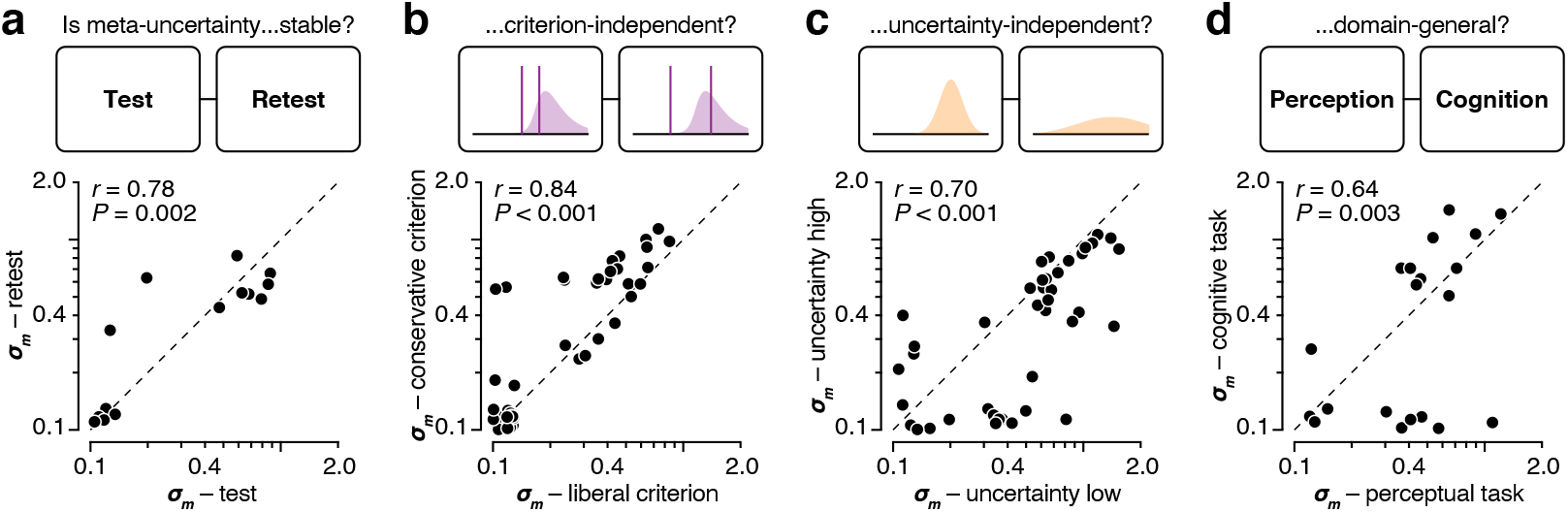
Evaluating meta-uncertainty as a psychological construct. (**a**) Comparison of meta-uncertainty estimates for 14 subjects who performed the same perceptual confidence task on two different occasions, separated by 1 month. We added a small amount of jitter to get a better view of overlapping data points in the lower left region of the plot. Meta-uncertainty was limited to a minimum value of 0.1. (**b**) Comparison of meta-uncertainty estimates for 43 sessions (performed by 32 subjects, see methods) whose 6-point confidence ratings were mapped onto a liberal and conservative 4-point rating scale. (**c**) Comparison of meta-uncertainty estimates for 43 sessions where subjects performed a confidence task involving low and high levels of stimulus uncertainty. (**d**) Comparison of meta-uncertainty estimates for 20 subjects who performed a perceptual and cognitive confidence task.

Under the CASANDRE model, meta-uncertainty provides a measure of metacognitive ability, not of confidence reporting strategy. To investigate whether this idealized pattern holds true in human choice-confidence data, we analyzed data from 43 sessions where subjects either performed a perceptual or a cognitive confidence task. They reported their confidence in a binary decision using a six-point rating scale ^24^. We artificially biased these confidence reports by mapping them onto a liberal and a conservative 4-point rating scale (see Methods) ^40^. This manipulation resulted in a mean confidence level of 2.89 and 2.43 – a substantial difference in light of the standard deviation (the effect size, expressed as Cohen’s *d*, is 3.16). We then used the model to analyze both perturbed versions of the data (see Methods). Meta-uncertainty estimates were strongly correlated (*r* = 0.84, *P* < 0.001; Fig. 5b), though note that they were on average higher for the conservatively biased version of the data (mean increase: 47%, median increase: 0%, *P* = 0.002, Wilcoxon signed rank test). This suggests that meta-uncertainty estimates are largely, but not fully, independent of subjects’ confidence reporting strategy.

We wondered whether meta-uncertainty depends on the absolute level of stimulus uncertainty ^41^. We analyzed data from 43 sessions where subjects either performed a perceptual or cognitive confidence task. In both tasks, stimulus uncertainty was manipulated by varying the variance of the category distributions over four levels ^24^. We used the CASANDRE model to analyze these data and estimated meta-uncertainty separately for the two lowest and the two highest levels of stimulus variance (see Methods). The former conditions resulted in a much higher task performance than the latter (average proportion correct decisions: 87% vs 70%). According to the model, the corresponding underlying levels of stimulus uncertainty, *σ*_*d*_, averaged 2.61 and 8.71. While increasing stimulus variance tripled stimulus uncertainty, meta-uncertainty estimates did not change much (median change: –14.76%, *P* = 0.004, Wilcoxon signed rank test). Moreover, meta-uncertainty estimates were strongly correlated across both sets of conditions (*r* = 0.70, *P* < 0.001; Fig. 5c). This suggests that meta-uncertainty is largely, but not fully, independent of the absolute level of stimulus uncertainty.

Whether metacognitive ability is domain-specific or domain-general is a debated question ^8,42–44^. We analyzed data from 20 subjects who performed a perceptual and cognitive confidence task. Both tasks had the same experimental design. Stimulus categories were either defined by the average orientation of a series of rapidly presented gratings, or by the average value of a series of rapidly presented numbers ^24^. Subjects’ performance level was correlated across both tasks (condition-specific proportion correct choices: *r* = 0.69, *P* < 0.001), and so were their reported confidence levels, albeit to a lesser degree (*r* = 0.53, *P* < 0.001). We used the CASANDRE model to analyze both data-sets (see Methods). Meta-uncertainty estimates were strongly correlated (*r* = 0.64, *P* = 0.003; Fig. 5d). Thus, meta-uncertainty appears to capture an aspect of confidence-reporting behavior that generalizes across at least some domains.

### Comparison with other metrics for metacognitive ability

Our method to analyze choice-confidence data is built on the hypothesis that metacognitive ability is determined by meta-uncertainty. It is natural to ask how this metric of metacognitive ability relates to alternatives. We approach this question in two ways. First, by investigating this relationship in silico whilst using the CASANDRE model as generative model of choice-confidence reports. And second, by comparing performance of these different candidate-metrics on a set of real bench-marking experiments (the tests shown in Fig. 5a-d).

One historically popular approach to quantify metacognitive ability consists of measuring the trial-by-trial correlation between choice accuracy and the confidence report (this metric is sometimes termed “phi”) ^12^. Consider an analysis of the choice-confidence reports of five model subjects who differed in their level of meta-uncertainty. We additionally varied the other model components in a step-wise fashion and computed phi for each simulated experiment. This analysis revealed a complex interdependence of the effects of the different model components on phi (Fig. 6a, top). An alternative method to quantify metacognitive ability that has gained popularity in recent years seeks to estimate how well confidence judgements distinguish correct from incorrect decisions ^13,45^. This estimate is expressed in signal-to-noise units and often termed “meta-*d′*“. The ratio of meta-*d* and stimulus discrimination ability (*d′*) theoretically provides a measure of metacognitive efficiency and is often considered the quantity of interest ^45^. Under the CASANDRE model, the meta-*d′*/*d′* metric does not provide a direct measurement of meta-uncertainty, but instead reflects a complex mixture of model components (Fig. 6a, middle).

**Figure 6.**
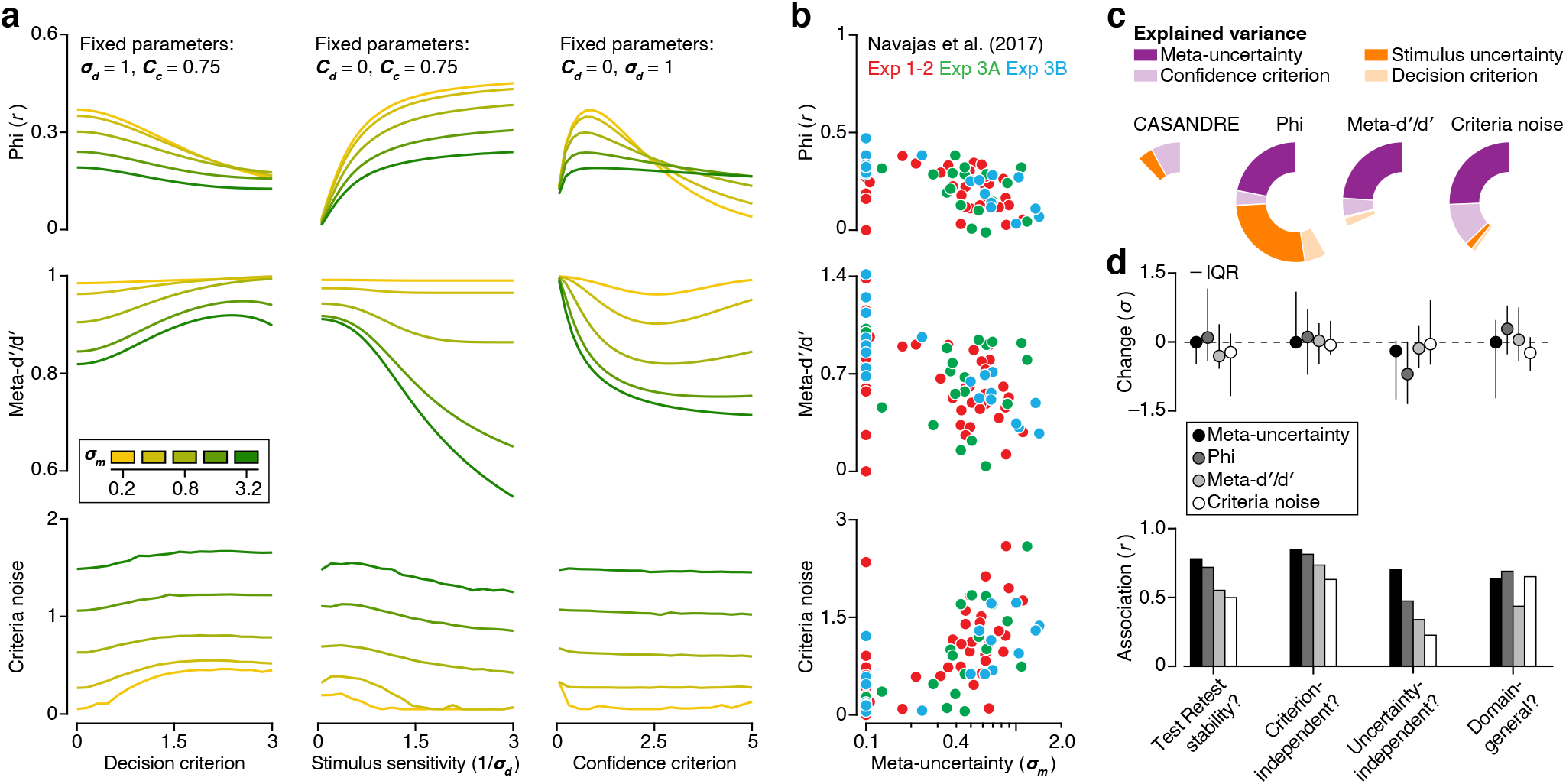
Comparing meta-uncertainty with three existing metrics of metacognitive ability. (**a**) We simulated choice-confidence data for a set of model observers who differed in their level of meta-uncertainty (colored lines) and additionally varied the decision criterion (left), the level of stimulus uncertainty (middle), and the confidence criterion (right). We estimated phi (top), meta-*d*′/*d*′ (middle), and criteria noise (bottom) for each simulated experiment. (**b**) Phi (top), meta-*d*′/*d*′ (middle), and criteria noise (bottom) plotted against meta-uncertainty estimates for three confidence experiments. Each symbol summarizes data from a single session (84 total sessions across 50 subjects, see methods). Meta-uncertainty was limited to a minimum value of 0.1. (**c**) Wedges indicate the proportion of variance in meta-uncertainty (left), phi, meta-*d*′/*d*′, *and criteria noise* explained by each model component. (**d**) Comparison of the performance of four metrics of metacognitive ability in four bench-marking tests. Top: analysis of estimation bias. Bottom: analysis of estimation robustness. Error bars illustrate the interquartile range (IQR) across subjects.

Finally, a recently introduced model of confidence judgments attributes metacognitive inefficiencies to perfectly correlated cross-trial variability in the confidence criteria ^28^. For experiments involving only two levels of stimulus strength, criteria noise can be estimated by fitting this model to choice-confidence data ^28^. In a practical sense, correlated criteria noise resembles meta-uncertainty in that it solely impacts confidence reports. However, assuming noisy uncertainty estimates versus noisy confidence criteria results in metrics that behave somewhat differently (Fig. 6a, bottom).

Now consider the relationship between these metrics and meta-uncertainty for the three experiments performed by Navajas et al. (2017). Meta-uncertainty estimates and phi are well correlated (*r* = –0.60, *P* < 0.001, Spearman correlation, Fig. 6b, top). But the correlation of two other model components with phi also reaches statistical significance: stimulus uncertainty (*r* = –0.59, *P* < 0.001) and the confidence criterion (*r* = –0.24, *P* = 0.028). Likewise, meta-uncertainty and meta-*d*′ /*d*′ are well correlated (*r* = –0.52, *P* < 0.001, Fig. 6b, middle). But the confidence criterion is also correlated with meta-*d*^*′*^/*d*′ (*r* = –0.29, *P* = 0.008).

Finally, meta-uncertainty and criteria noise are well correlated (*r* = 0.64, *P* < 0.001, Fig. 6b, bottom). But the confidence criterion is also correlated with criteria noise (*r* = 0.37, *P* < 0.001). Variability in each of these metrics of metacognitive ability thus in part reflects variability in meta-uncertainty, and in part variability in other components of the CASANDRE model.

To identify the relative importance of the different model components, we decomposed the variance of these metrics using the averaging-over-orderings technique (see Methods) ^46,47^. We first asked whether variability in meta-uncertainty could be explained by other model components, but found this not be be the case (fraction of explained variance: 13%, Fig. 6c). In contrast, variability in phi is predominantly explained by stimulus uncertainty (27%), followed by meta-uncertainty (22%). For meta-*d*′ /*d*′ and criteria noise, most variance is explained by meta-uncertainty (24% and 26%) while the contribution of the other model components is rather small (Fig. 6c). In summary, for all three alternative metrics, about three quarters of the variance arises from factors other than meta-uncertainty.

Our analysis suggest that phi, meta-*d*′ /*d*′, and criteria noise do not isolate the factors that limit metacognitive ability but instead measure a complex mixture of factors underlying choice-confidence data. We wondered how the performance of these mixtures in bench-marking experiments compares to that of meta-uncertainty. We computed phi, meta-*d*′ /*d*′, and criteria noise for the data sets shown in Fig. 5a-d. For each test, we first asked whether the manipulation induced a systematic change in the range of the different metrics. This was generally not the case. Change, expressed in units of standard deviation, tended to be small for all four metrics (Fig. 6d, top). We then asked for each test whether the different metrics were correlated across both test conditions. Correlations ranged from weak to strong levels, with three tests failing to reach statistical significance (uncertainty independence of criteria noise: *r* = 0.23, *P* = 0.145; test-retest reliability of criteria noise: *r* = 0.50, *P* = 0.072; and domain generality of meta-*d*′ /*d*′ : *r* = 0.44, *P* = 0.056). Overall, meta-uncertainty compared favorably to the alternative metrics. The mean correlation value across the four tests was 0.74 for meta-uncertainty, 0.67 for phi, 0.52 for meta-*d*′ /*d*′, and 0.50 for criteria noise (all correlations are shown in Fig. 6d, bottom).

### Manipulating meta-uncertainty

Can metacognitive ability be manipulated experimentally? Key to our framework is that confidence judgements require a subject to estimate uncertainty on a trial-by-trial basis. This becomes more difficult when experiments involve more confusable levels of stimulus uncertainty. We therefore expect that meta-uncertainty will grow with the number of stimulus uncertainty levels. To appreciate our logic, consider the ideal Bayesian uncertainty estimation strategy which consists of combining information obtained from ambiguous sensory measurements with prior task-specific knowledge. Specifically, the sensory measurement informs the uncertainty likelihood function, while knowledge of task statistics (i.e., the distribution of stimulus uncertainty levels) is summarized in a prior uncertainty belief function (Fig. 7a). The combination of both yields a posterior uncertainty belief function, the maximum of which is the “best possible” uncertainty estimate (Fig. 7a). Due to noise, repeated presentations of the same condition will yield different likelihood functions (Fig. 7a, see Methods). If the task involves only one level of stimulus uncertainty, the prior is a fixed delta function, and so is the posterior. Consequently, the maximum posterior estimate will not vary across trials and the ideal estimation strategy results in zero meta-uncertainty. However, when a task involves multiple levels of stimulus uncertainty, the prior will be more dispersed, causing the resulting maximum posterior estimate to be more variable across trials. Under an ideal Bayesian estimation strategy, meta-uncertainty thus initially grows with the number of uncertainty levels (Fig. 7b). We wondered whether this normative prediction affords insight into human metacognition. To test this hypothesis, we used the CASANDRE model to analyze six confidence experiments that varied in the number of randomly interleaved uncertainty levels (see Methods). These experiments utilized different stimuli and employed different experimental designs ^24–28^. Yet, as expected, meta-uncertainty appears to grow lawfully with the number of uncertainty levels (Fig. 7e).

**Figure 7.**
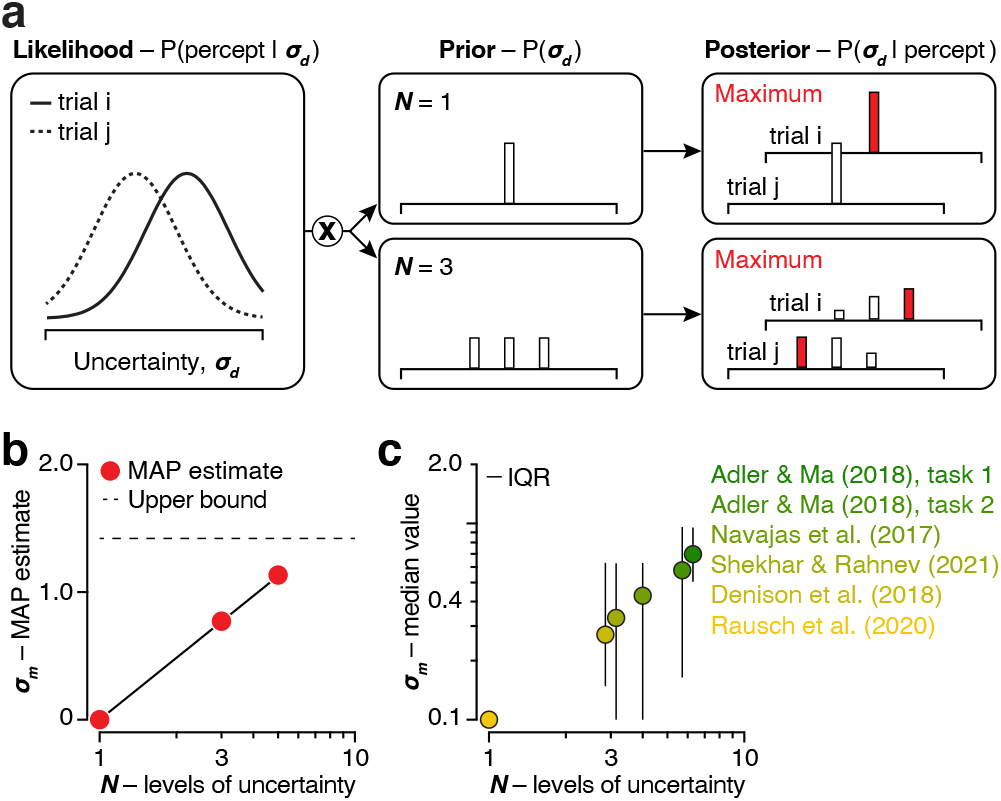
Meta-uncertainty depends on task structure. (**a**) We studied how meta-uncertainty depends on the number of uncertainty levels under an ideal Bayesian uncertainty estimation strategy. The likelihood of each uncertainty value is computed from a sensory measurement (left) while a prior belief function specifies task-specific knowledge of possible uncertainty values (middle). The product of the prior and likelihood gives the posterior (right). Due to noise, the likelihood function will differ across repeated trials (left: full vs dotted line). The impact of this variability on the posterior depends on the dispersion of the prior (right: top vs bottom panel). (**b**) Meta-uncertainty plotted against number of uncertainty levels for the ideal Bayesian estimator. The upper bound (dotted line) is set by the cross-trial variability of the maximum of the likelihood function and is reached when the prior is a uniform distribution. (**c**) Median level of meta-uncertainty plotted against number of uncertainty levels for six confidence experiments. Meta-uncertainty was limited to a minimum value of 0.1. Error bars illustrate the interquartile range (IQR) across subjects (Adler and Ma (2018) task 1: 19 subjects; task 2: 34 subjects; Navajas et al. (2017): 50 subjects; Shekhar and Rahnev (2021): 20 subjects; Denison et al. (2018): 12 subjects; Rausch et al. (2020): 25 subjects).

## Discussion

It has long been known that humans and other animals can meaningfully introspect about the quality of their decisions and actions ^5–7,31,48^. Quantifying this ability has remained a significant challenge, even for simple binary decision-making tasks ^12,13,15,28,40,41^. The core problem is that observable choice-confidence data reflect metacognitive ability as well as task difficulty and response bias. To overcome this problem, we introduced a metric that is anchored in an explicit hypothesis about the decision-making process that underlies behavioral reports. Our method is based on likening choice-confidence data to the outcome of an abstract mathematical process in which confidence reflects a subject’s noisy estimate of their choice reliability, expressed in signal-to-noise units ^14,20,49^. This framework allowed us to specify the effects of factors that limit metacognitive ability and to summarize this loss in a single, interpretable parameter: meta-uncertainty. We showed that this process model (which we term the CASANDRE model) can explain the effects of stimulus strength and stimulus reliability on confidence reports and that meta-uncertainty can be estimated from conventional experimental designs. We found that a subject’s level of meta-uncertainty is stable over time and across at least some domains. Meta-uncertainty can be manipulated experimentally: it is higher in tasks that involve more levels of stimulus reliability. Meta-uncertainty appears to be mostly independent of task difficulty and confidence reporting strategy. Widely used metrics for metacognitive ability are poor proxies for meta-uncertainty. As such, the CASANDRE model represents a notable advance toward realizing crucial medium and long-term goals in the field of metacognition ^50^.

The mental operations underlying confidence in a decision have long intrigued psychologists. Two key unresolved issues are the structure and nature of the confidence computation ^50^. At stake are two intertwined questions: (1) Does confidence arise from a single, dual, or hierarchical process? and (2) What exactly does confidence reflect? Some authors have proposed that decision outcome and confidence both arise from a single stimulus strength estimation process ^31,51–53^. Such models can explain the effects of stimulus strength, but not of stimulus reliability. Others have argued in favor of a dual process in which decision outcome and confidence are based on different stimulus strength estimates ^54–56,56,57^. This may be the appropriate framework for cases in which subjects acquire additional task-relevant information after reporting their choice ^57–60^. For all other cases, it appears overly complex. Instead, we have modeled confidence judgements as arising from a hierarchical process ^61^. The first stage determines the choice, the second stage determines confidence (Fig. 1e). We found that this model structure systematically outperforms a single stage alternative (Fig. 3c, top). The structure of the computation clarifies its nature. Many previous studies are built on the premise that confidence reflects a subject’s assessment of decision accuracy (“What is the probability that my choice is correct?”). This premise directly motivates Bayesian models of confidence ^1,25,31,62–68^ and tacitly underlies popular metrics of metacognitive ability ^13,20^. However, when experimental manipulations bias perceptual choices, aggregated confidence reports do not track choice accuracy but choice consistency ^20,36,37^. At the single trial level, this suggests that confidence reflects a subject’s assessment of decision reliability (“What is the probability that I would make the same choice again?”, see equation 1). For an unbiased subject who is choosing between two alternatives, decision accuracy and decision reliability are indistinguishable ^20,67^. Yet, the distinction matters greatly, as it implies that the same computation that underlies confidence in decisions with a well-defined correct and incorrect option may generalize to subjective domains that lack this feature (“Which political candidate will I support?”, “Which beer will I have?”, “Should I skip class today?”). ^69^

Key to our proposal is that assessing the reliability of a decision requires the use of additional information (stimulus uncertainty) ^21^ that in most tasks has no relevance for the choice as such. The notion that subjects can incorporate a stimulus uncertainty estimate when making perceptual inferences is well established ^25,70–72^. And there is considerable evidence that neural activity in sensory areas of the brain conveys information about stimulus features as well as the uncertainty of those features ^68,73–78^. Our proposed confidence computation yielded a new prediction: the more levels of stimulus uncertainty a task involves, the more variable uncertainty estimates will be. We validated this prediction by analyzing data from six different confidence experiments in which 160 subjects completed a total of 243,000 trials (Fig. 7c). This finding is arguably the strongest piece of empirical evidence that meta-uncertainty is the critical factor that limits human metacognitive ability. It was enabled by the use of modern computational tools to quickly compute the approximate ratio of two distributions (i.e., the confidence variable distribution) and by the availability of the confidence database ^23^. This phenomenon also raises the question to what degree metacognitive ability estimates are influenced by experimental design. For example, studies that increase the volatility of stimuli within a trial (thereby making uncertainty more difficult to estimate) report confidence distortions that could likely be captured by the CASANDRE model ^79–81^. An important future direction will be to investigate the effect of different stimulus uncertainty distributions on metacognitive ability.

The CASANDRE model provides a static description of the outcome of a hierarchical decision-making process. However, making a decision requires time. The more difficult the decision, the more time it requires ^82,83^. For this reason, some authors have suggested that decision time directly informs confidence ^58,84^. This proposal enjoys strong empirical support ^38,63,80,84^. It is related to our proposed confidence computation, provided that the decision variable results from a mechanism that resembles bounded evidence accumulation ^7^. For these mechanisms, time to reach the bound reflects the drift rate of a drift diffusion process. Drift rate is governed by stimulus strength, normalized by stimulus uncertainty and thus determines decision reliability. Moreover, just like our second stage involves an additional factor to reflect on the quality of the decision (the uncertainty estimate), time measurements are not inherent to bounded accumulation. Like uncertainty estimates, neural and behavioral time measurements are strictly positive and noisy ^85–87^. This noisiness provides the conceptual dynamic analogue for meta-uncertainty in our static model. How it affects confidence reporting behavior in the diffusion-to-bound framework has not yet been studied. It remains to be seen whether choice outcome, reaction time, and metacognitive ability can all be modeled simultaneously.

Process models are powerful tools to study cognition and perception. Here we leveraged a process model to interrogate the computations underlying our sense of confidence, to determine the effectiveness of various experimental designs, and to examine model recoverability. However, the usefulness of process models far exceeds our current application. Specifically, when coupled to an explicit goal such as maximizing choice accuracy, process models can be used to derive the optimal task strategy. The resulting predictions offer a critical point of reference for human behavior ^88^. This approach has revealed that humans improve the quality of uncertain decisions by accumulating evidence over time ^82^, by combining information acquired through different sensory modalities ^70^, and by exploiting knowledge of statistical regularities in the environment ^89^. Might the same hold true for uncertain confidence judgments? Stated more generally: Does our brain attempt to maximize the precision of our sense of confidence? This is a fundamental question that is ripe to be addressed. Doing so will require experiments that manipulate meta-uncertainty and incentivize the confidence reporting strategy (e.g., refs. ^31,48,53,55,63,90–92^). The process model we have developed provides a vehicle to derive the reward-maximizing strategy and to evaluate whether human meta-uncertainty changes as expected for theoretically ideal introspection. We took a first step in this direction and validated a novel prediction: meta-uncertainty changes with task-structure as expected under an ideal Bayesian uncertainty estimation strategy.

## Methods

### Modeling: Hierarchical decision-making process

We model choice-confidence data in binary decision-making tasks as arising from a hierarchical process. The first stage follows conventional signal detection theory applications ^22^ and describes the primary decision as resulting from the comparison of a one-dimensional decision variable, *V*_*d*_, with a fixed criterion, *C*_*d*_. The decision variable is subject to zero-mean Gaussian noise and hence follows a normal distribution with mean *μ*_*d*_ and standard deviation *σ*_*d*_. The decision variable is converted into a signed confidence variable, 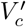, by taking the difference of *V*_*d*_ and *C*_*d*_, and dividing this difference by 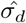, the subject’s estimate of *σ*_*d*_. The family of normal distributions is closed under linear transformations. This means that, if 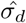 were a constant, 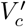 would also follow a normal distribution with mean 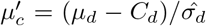 and standard deviation 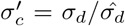. The confidence report results from the comparison of the confidence variable with a single fixed criterion, *C*_*c*_ (or with a set of criteria if the confidence scale has more than two levels). It follows that the probability of a “confident” judgement given a “Category A” decision is given by *P* (*C* = 1 | *D* = 0) = Φ(*C*_*c*_), where Φ (.) is the cumulative normal distribution with mean 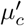 and standard deviation 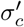. By the same logic, *P* (*C* = 0 | *D* = 0) = Φ (0) − Φ (− *C*_*c*_), *P* (*C* = 0 | *D* = 1) = Φ (*C*_*c*_) − Φ (0), and *P* (*C* = 1 | *D* = 1) = 1 − Φ (*C*_*c*_). Key to the CASANDRE model is that 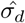 is not a constant, but a random variable that follows a log-normal distribution with mean *σ*_*d*_ and standard deviation *σ*_*m*_. Consequently, the signed confidence variable is a mixture of normal distributions, with mixing weights determined by *σ*_*m*_. To obtain the probability of each response option under this mixture, we sample 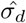 in steps of constant cumulative density (using the Matlab function ‘logninv’), compute the probability of each response option under each sample’s resulting normal distribution (using the Matlab function ‘normcdf’), and average these probabilities across all samples. We found that this procedure yields stable probability estimates once the number of samples exceeds 25 (i.e., sampling the log-normal distribution in steps no greater than 4%). For all applications in this paper, we used 100 samples, thus sampling 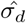 at a cumulative density of 0.5%, 1.5%, 2.5%,…, and 99.5%. Finally, note that whenever we report values for *σ*_*m*_, we use the coefficient of variation (*σ*_*m*_/*σ*_*d*_), as this ratio is identifiable under the model (the absolute level of meta-uncertainty is not, just like the absolute level of *σ*_*d*_ cannot be uniquely estimated from choice data).

### Modeling: Parameterization, simulations, and fitting

We analyzed data from a large set of previously published studies that employed different task designs. The simplest designs involve the combination of a 2-AFC categorization decision and a binary confidence report (i.e. the model simulations shown in Fig. 2 and 4). Under the CASANDRE model, the predicted probability of each response option is fully specified by five parameters: the mean of the decision variable (*μ*_*d*_), the standard deviation of the decision variable (*σ*_*d*_), the decision criterion (*C*_*d*_), the level of meta-uncertainty (*σ*_*m*_), and the confidence criterion (*C*_*c*_). It is not possible to estimate each of these parameters for every unique experimental condition. To make the model identifiable, we generally assume that *μ*_*d*_ is identical to the true stimulus value, that *σ*_*d*_ is constant for a given level of stimulus reliability, and that *C*_*d*_, *σ*_*m*_ and *C*_*c*_ are constant across multiple conditions.We limited *σ*_*m*_ to a minimum value of 0.1, as values below this had indistinguishable effects on model behavior. Figure 2 shows how each of the parameters affects the model’s behavior. Finally, when fitting data, we use one additional parameter, *λ*, to account for stimulus-independent lapses ^93^, which we assume to be uniformly distributed across all response options. We fit the model on a subject-by-subject basis. For each subject, we compute the log-likelihood of a given set of model parameters across all choice-confidence reports and use an iterative procedure to identify the most likely set of parameter values (specifically, the interior point algorithm used by the Matlab function ‘fmincon’). Figure 4b shows an example model fit to a synthetic data set whereby we used 5 free parameters (*λ, σ*_*d*_, *C*_*d*_, *σ*_*m*_, and *C*_*c*_) to capture data across 20 experimental conditions.

Some studies used a task design that combined a 2-AFC categorization decision with a multi-level confidence rating scale (i.e., ref. ^24,25,27,28^). To model these data, we used the same approach as described above but we used multiple confidence criteria (one less than the number of confidence levels). We modeled the data from ref. ^27^ using seven free parameters: *λ, σ*_*d*_, *C*_*d*_, *σ*_*m*_, and *C*_*c*_ (4-point confidence rating scale, thus three in total) (see Fig. 7c and Supplementary Fig. 5a). We modeled some data from ref. ^25^ (task 1) using seventeen free parameters: *λ, σ*_*d*_ (one per contrast level, six in total), *C*_*d*_ (one per contrast level, six in total), *σ*_*m*_, and *C*_*c*_ (4-point confidence rating scale, thus three in total). Example fits are shown in Fig. 1b,c and in Supplementary Fig. 1 (also see Fig. 7c, task 1 and Supplementary Fig. 5e). We modeled the data from ref. ^24^ using twelve free parameters: *λ, σ*_*d*_ (one per stimulus variance level, four in total), *C*_*d*_, *σ*_*m*_, and *C*_*c*_ (6-point confidence rating scale, thus five in total). Example fits are shown in Supplementary Fig. 4 (also see Fig. 7c, Fig. 6b-d, and Supplementary Fig. 5d). We modeled the data from ref. ^28^ using 10 free parameters: *σ*_*d*_ (one per stimulus reliability level, three in total), *C*_*d*_, *σ*_*m*_, and *C*_*c*_ (continuous confidence rating scale, discretized into 6-point confidence rating scale, thus five in total). See Fig. 7c, and Supplementary Fig. 5b).

Some studies used a task design in which the 2-AFC categorization decision pertained to two category distributions with the same mean but different spread (i.e., ref. ^25,26^). To model these data, we assumed that the primary decision results from a comparison of the decision variable with two decision criteria, and that the confidence estimate is based on the distance between the decision variable and the nearest decision criterion. We modeled some data from ref. ^25^ (task 2) using twenty-three free parameters: *λ, σ*_*d*_ (one per contrast level, six in total), *C*_*d*_ (two per contrast level, twelve in total), *σ*_*m*_, and *C*_*c*_ (4-point confidence rating scale, thus three in total). See Fig. 7c, task 2. Example fits are shown in Supplementary Fig. 6 (also see Supplementary Fig. 5e). We modeled data from ref. ^26^ using twenty-two free parameters: *λ, σ*_*d*_ (one per attention level, three in total), *C*_*d*_ (two per attention level, six in total), *σ*_*m*_ (one per attention level, three in total), and *C*_*c*_ (4-point confidence rating scale, one set per attention level, thus nine in total). See Fig. 7c and Supplementary Fig. 5c.

Some studies used a task design that combined a 2-IFC categorization decision with a confidence report (i.e., Arbuzova and Filevich, unpublished but available in the Confidence Database ^23^). In these tasks, a subject is shown two stimulus intervals and judges which interval contained the “signal” stimulus. To model such data, we assume that the decision is based on a comparison of the evidence provided by each stimulus interval. The one-dimensional decision variable, *V*_*d*_, reflects the outcome of this comparison, which we model as a difference operation ^22^. The difference of two Gaussian distributions is itself a Gaussian with mean equal to the difference of the means and standard deviation equal to the square root of the sum of the variances. Everything else is the same as for the 2-AFC task. When different from zero, *C*_*d*_ now reflects an interval bias (e.g., a preference for “interval 1” choices). See example fit in Fig. 1d.

### Modeling: Model comparison

We evaluated CASANDRE’s assumed confidence computation and overall model architecture by fitting different model variants to an experiment that involved joint manipulations of stimulus strength and stimulus reliability (ref. ^25^, task 1, 19 subjects). For each model comparison, we computed the Akaike Information Criterion, given by:

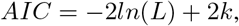

where *L* is the maximum value of a model’s likelihood function and *k* is the number of fitted parameters. To focus this analysis on the model’s second-stage operations, we began by fitting 13 first-stage parameters to each subject’s choice data: *λ, σ*_*d*_ (one per contrast level, six in total), *C*_*d*_ (one per contrast level, six in total). These parameters were kept constant across all model variants. The head-to-head model comparisons consisted of (1) confidence as a noiseless stimulus strength estimate vs confidence as a noiseless decision reliability estimate, (2) confidence as a noiseless decision reliability estimate vs confidence as a noisy decision reliability estimate, (3) symmetric confidence criteria vs asymmetric confidence criteria, and (4) a log-normal vs Gamma second-stage noise distribution.

### Datasets

The majority of our analyses focus on two studies ^24,25^. To test the effect of task structure on meta-uncertainty, we additionally analyzed data from three other studies ^26–28^. The data from Navajas et al. (2017) were provided by an author ^24^. All other datasets were obtained from the Confidence Database ^23^ (available at: https://osf.io/s46pr/). Given that the CASANDRE model yields more reliable parameter estimates for longer experiments with more stimulus conditions (error bars in Fig. 4c), we included all experiments from the database that involved a large number of subjects, several hundred trials per subject, and multiple levels of stimulus strength and/or stimulus reliability. All detailed experimental designs and procedures are available in the original publications or in abbreviated form in the Confidence Database. We briefly describe each data set below.

We analyzed data from all three experiments in ref. ^25^. All subjects in experiments 1 and 2 performed both task 1 (discriminating categories of orientation distributions with different means but the same standard deviation; their “Task A”) and task 2 (discriminating categories of orientation distributions with the same mean but different standard deviations; their “Task B”). Since stimulus orientations were drawn from a continuous distribution, to plot the data we grouped nearby orientations into 9 bins with similar numbers of trials. Data and model fits from two example subjects performing task 1 in experiment 1 are shown in Fig. 1b-c and Supplementary Fig. 1. Fitted parameters from all 19 subjects who performed experiments 1 and 2 are included in Fig. 7c (task 1) and Supplementary Fig. 5f. Subjects in experiment 3 performed only task 2. Data and model fits from an example subject performing task 2 in experiment 3 are shown in Supplementary Fig. 6. Fitted parameters from all 34 subjects who performed task 2 in experiments 1, 2, and 3 are included in Fig. 7c (task 2) and Supplementary Fig. 5e.

We analyzed data from all 3 experiments in ref. ^24^. 30 subjects performed experiment 1. 14 of those 30 subjects returned about a month after their first session to perform the same task again as experiment 2. Finally, 20 subjects performed experiment 3, participating in a perceptual (experiment 3A) and cognitive (experiment 3B) task in two different sessions. We analyzed each of these 84 different experimental sessions independently. Data and model fits from an example subject are shown in Supplementary Fig. 4. Fitted parameters and alternative metacognitive metrics from all 14 subjects who performed both experiments 1 and 2 are included in Fig. 5a and Fig. 6d (Test-retest stability). Fitted parameters and alternative metacognitive metrics from all 20 subjects who performed experiment 3 are included in Fig. 5d and Fig. 6d (Domain generality). Fitted parameters from 50 subjects performing experiment 1 and the perceptual task of experiment 3 (experiment 3A) are included in Fig. 7c and Supplementary Fig. 5d. Further analyses using these data to test the independence of meta-uncertainty from confidence reporting strategy and uncertainty are explained in the next section.

We analyzed unpublished data from Arbuzova and Filevich (available in the Confidence Database under the name Arbu-zova_unpub_1) ^23^. This experiment demonstrates the generalization of the CASANDRE model to a visuomotor estimation task as well as 2-IFC experimental designs. Data and model fits from a representative subject are shown in Fig. 1d.

Fitted parameters from all 25 subjects from ref. ^27^ and from all 20 subjects from ref ^28^ are included in Fig. 7c. We analyzed data from 12 subjects performing a version of task 2 in ref. ^25^ with an added attention manipulation from ref. ^26^. To get the single estimate of meta-uncertainty included in Fig. 7c for each subject, we averaged the values estimated from all three attention conditions, as these were not significantly different.

### Construct validity analyses

To test the independence between confidence reporting strategy and measures of metacognitive ability, we manipulated the confidence reporting behavior of subjects across all sessions from ref. ^24^ (following an analysis developed by ref. ^40^). In these experiments, confidence reports were measured using a six-point rating scale. We remapped responses into a four point rating scale using two different grouping rules (one conservative, one liberal). The conservative mapping is [1|2 3 4|5|6], the liberal mapping is [1|2|3 4 5|6] (i.e., for the conservative mapping, ratings 2, 3 and 4 were combined, and for the liberal mapping, ratings 3, 4, and 5 were combined.) To limit the model comparison to the second stage of the decision making process, the lapse rate, stimulus sensitivity, and perceptual criterion were shared across both model variants. Only the meta-uncertainty and confidence criteria differed across both model variants. To obtain adequately constrained and stable model fits to these manipulated data, we only included a session in the analysis if at least 10 responses were recorded at the highest level of the confidence scale. This reduced a total of 84 sessions to 43 (and 50 subjects to 32), shown in Fig. 5b.

To test the independence between stimulus uncertainty and measures of metacognitive ability, we split experimental data from each session in half ^24^. We estimated meta-uncertainty independently for the two easiest and the two hardest stimulus conditions. To limit the model comparison to the question of whether meta-uncertainty is independent of stimulus reliability, all other model parameters were fixed across conditions. For consistency with the criterion analysis, we applied the same inclusion criteria, yielding data from 43 sessions included in Fig. 5c.

### Calculating alternative metacognitive metrics

We probed the relation between meta-uncertainty and three alternative metrics of metacognitive ability under the CASANDRE model. We used two distinct procedures for this. First, to obtain estimates of “meta-*d*′ “, we used the CASANDRE model to specify the probability of each response option in a 2-AFC discrimination task with binary confidence report options for an experiment that included two stimulus conditions. We calculated meta-*d*′ following ref. ^45^. Briefly, we searched for the level of sensory noise and the confidence criterion that best explained the distribution of confidence reports conditioned on the primary choice, assuming a normally distributed confidence variable. The ratio of the ground truth sensory noise level and this estimate is plotted in the middle panels of Fig. 6a. Second, to obtain the expected value of phi, we simulated 200,000 trials in an experiment that included 20 levels of stimulus strength. We then calculated the Pearson correlation between the resulting choice accuracy and confidence vectors (Fig 6a, top panels). We used these same simulated trials to fit the criteria-noise model of Shekhar and Rahnev ^28^. We downloaded their parameter optimization code and modified it as appropriate to fit our simulated data (available at https://osf.io/s8fnb/). In their procedure, a nested two-step, coarse-to-fine search algorithm is used to optimize the estimated confidence criteria and the confidence criteria noise level. The resulting criteria noise estimates are plotted in the bottom panels of Fig 6a. The non-smooth appearance of the curves is a consequence of instabilities in the fitting procedure.

We also computed these alternative metrics for each session from ref. ^24^ (see Fig. 6b-d). As is conventional, we estimated *d*′ for each stimulus condition from the observed hit and false alarm rates ^22^. To obtain estimates of “meta-*d*′ “, we searched for the decision criterion, the set of confidence criteria, and the level of sensory noise that best explained the choice-conditioned data, assuming a normally distributed confidence variable. To obtain a single meta-*d*′ /*d*′ estimate per session, we computed the arithmetic mean across the four stimulus conditions. We computed phi for each session by calculating the Pearson correlation between choice accuracy and raw confidence report. We again used the fitting procedure of Shekhar and Rahnev ^28^, estimating decision criterion and four values of *d*′ and optimizing four sets of confidence criteria and the value of criteria noise across the four stimulus conditions (Fig. 6b-d).

To compute the proportion of variance in each alternative metric across 84 sessions ^24^ explained by different components of the CASANDRE model, we used the averaging-over-orderings technique ^46,47^. We used multiple linear regression to obtain the variance in a metric explained by the CASANDRE model. Then, for each model parameter we compute the difference in explained variance when the parameter is included and when it is not. The resulting estimates of explained variance for each parameter are plotted in Fig. 6c.

### Bayesian uncertainty estimation

We examined a simple model of Bayesian uncertainty estimation (Fig. 7a-b.). We modeled the uncertainty likelihood function as a Gaussian function with a mean value, *μ*_*u*_, that varied from trial-to-trial. Each trial, *μ*_*u*_ was randomly drawn from a Gaussian distribution whose average matched the true level of stimulus uncertainty, *S*_*u*_, and with standard deviation *σ*_*u*_. As is typical for a well-calibrated model, the spread of the likelihood function equalled *σ*_*u*_. We assumed three different experimental designs that yielded a prior uncertainty belief function composed of a single delta function (*N* = 1), three delta functions (*N* = 3), and five delta functions (*N* = 5). We simulated 1000 trials per design. In this simulation, we computed the posterior on a single trial basis and selected its maximum as the MAP uncertainty estimate. Fig 7b summarizes a simulation in which *S*_*u*_ = 2.5, *σ*_*u*_ = 1.5, and the prior belief function peaked at 2.5 for *N* = 1, at 1.67, 2.5, and 3.33 for *N* = 3, and at 0.83, 1.67, 2.5, 3.33, and 4.17 for *N* = 5.

## Data availability

This study generated no new data. The data used in this study are available from the Confidence Database (available at: https://osf.io/s46pr/).

## Code availability

The code supporting the findings of this study and a software package implementing the CASANDRE model is publicly available (https://github.com/gorislab/CASANDRE.git).

## Acknowledgements

We thank the creators and contributors to the Confidence Database and Joaquin Navajas for making their data available. This work was supported by a U.S. National Science Foundation Graduate Research Fellowship (to ZMB-S), U.S. National Institutes of Health grants T32 EY021462 (supporting CMZ), K99 EY032102 (to CMZ), and EY032999 (to RLTG), and a Whitehall Foundation grant (to RLTG).

## Author contributions

ZMB-S, CMZ, and RLTG conceived the study, developed the theory, performed the simulations, analyzed the data, and wrote the paper.

## Supplementary information

### Adler and Ma (2018), task 1

Figure 1b,c shows data from two subjects who performed a perceptual 2-AFC categorization task and additionally reported their confidence using a four-point rating scale. Data were collected by Adler and Ma (2018). Supplementary Figure 1a,b illustrates model fits by plotting the psychometric function (top row) and accompanying confidence function (bottom row) for each stimulus contrast (columns). Subjects completed 2,160 trials each. To model these data, we used one lapse rate parameter (obs 1-6: 5%; obs 1-9: 0.5%), one contrast-specific sensitivity parameter (obs 1-6: 0.21, 0.15, 0.14, 0.11, 0.07, and 0.06; obs 1-9: 0.55, 0.42, 0.36, 0.24, 0.16, and 0.10), one contrast-specific decision criterion parameter (obs 1-6: 0.62, 0.55, 0.00, 1.49, 0.45, and 1.28 degrees; obs 1-9: 0.03, -0.08, -0.32, 0.05, -0.35, and -1.33 degrees), one meta-uncertainty parameter (obs 1-6: 0.21; obs 1-9: 0.51), and three confidence criterion parameters (obs 1-6: 0.02, 0.40, and 1.92; obs 1-9: 1.42, 3.30, and 10.99). The log-probability of the data under the model was –3,462.1 for obs 1-6, and –2,654.0 for obs 1-9.

**Supplementary Figure 1.**
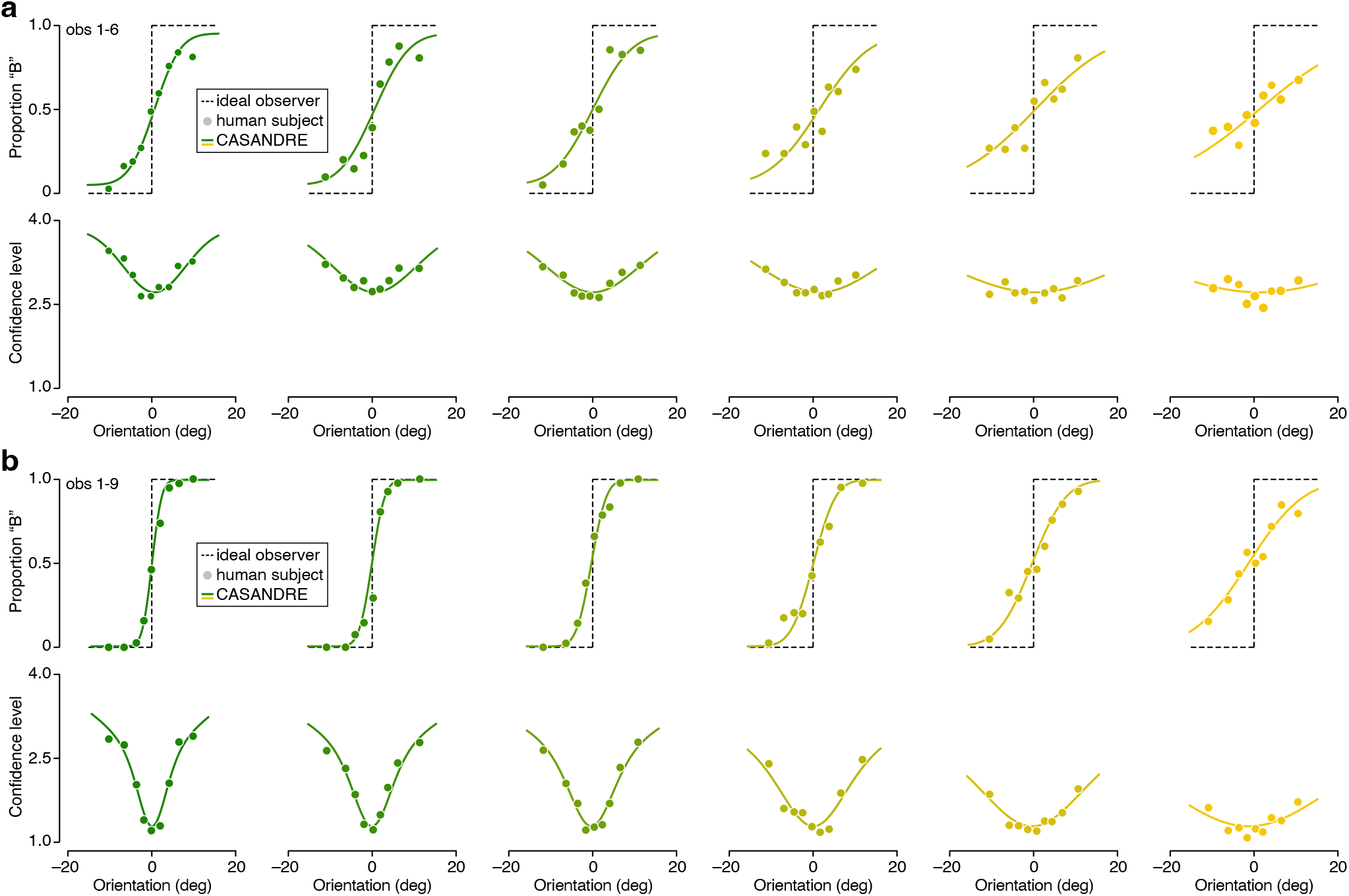
Model fits for two example subjects from Adler and Ma (2018). Both subjects judged whether a stimulus belonged to category A or B. Category A stimuli typically had an orientation smaller than zero, while category B stimuli typically had an orientation larger than zero. Stimuli varied in orientation and contrast. (**a**) Top: Proportion of “Category B” choices is plotted against stimulus orientation, split out by stimulus contrast (columns, contrast decreases from left to right), for one example subject (observer 6 in experiment 1 from ref. ^25^). Bottom: Same for mean confidence level. Symbols summarize observed choice behavior, the dotted line illustrates the theoretical optimum, and the full lines show the fit of a two-stage process model of decision-making. Symbol size is proportional to the number of trials. (**b**) Same for a different example subject (observer 9 in experiment 1 from ref. ^25^).

### Evaluating the ideal of criterion-symmetry

Figure 3a,b shows data for one example subject (2-7) from Adler and Ma (2018) fit with different model variants. This example subject’s data were marginally better fit with a model with asymmetrical rather than symmetric confidence criteria (AIC difference = 10.9). Supplementary Figure 2 illustrates data from another example subject (2-2) for whom the difference in model performance is more substantial (AIC difference = 149.2).

**Supplementary Figure 2.**
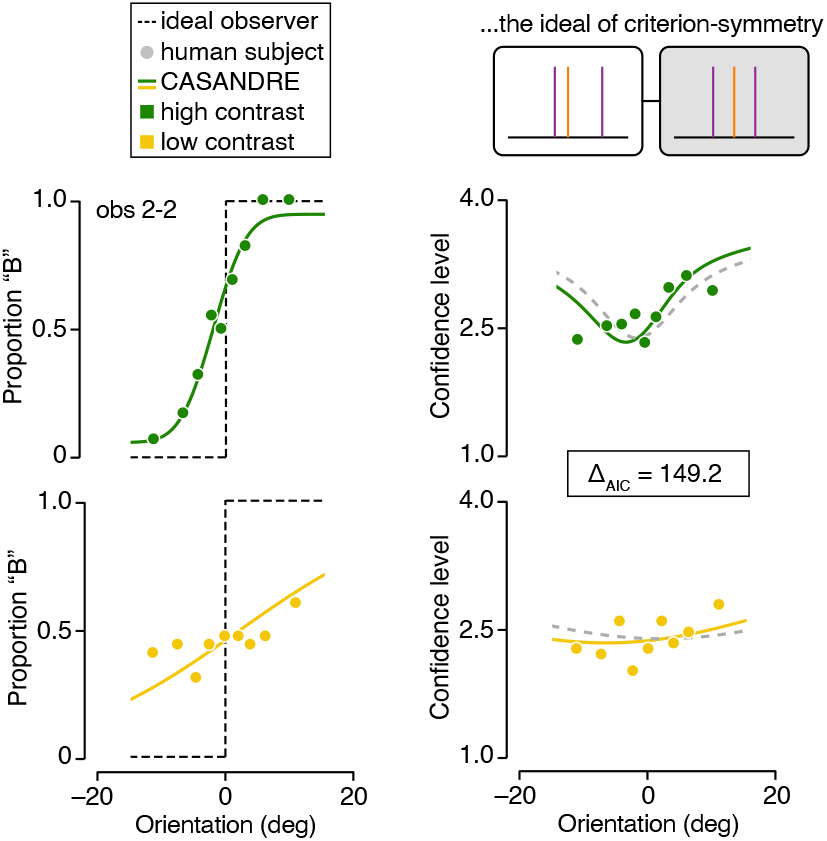
Following the same conventions as Figure 3. Left: proportion of “Category B” choices is plotted against stimulus orientation for high contrast (top, green) and low contrast (bottom, yellow) for example subject (observer 2 in experiment 2 from ref. ^25^). Right: Mean confidence level is plotted as a function of stimulus orientation for the same example observer. Symbols indicate data; lines indicate model fits. Solid lines indicate CASANDRE model fit with asymmetric confidence criteria. Dashed lines indicate CASANDRE model fit with symmetric confidence criteria.

### Meta-uncertainty recovery, Adler and Ma (2018) task 1

We verified that meta-uncertainty can be reliably recovered for the datasets used to evaluate the model’s architecture. These datasets came from the 19 subjects who participated in ref. ^25^‘s experiment 1 and 2. For each subject, we simulated 100 choice-confidence datasets using the CASANDRE model and the best-fitting parameter values. Each simulated experiment exactly matched the set of trials completed by the subject. We then analyzed the synthetic data in the same manner as the real data. As can be seen in Supplementary Figure 3, under this experimental design, meta-uncertainty is recoverable.

**Supplementary Figure 3.**
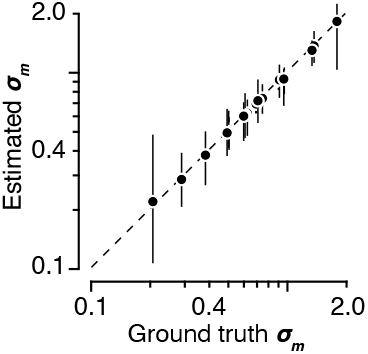
Model recovery analysis for Adler and Ma (2018) task 1 data. The median estimate of meta-uncertainty is plotted against the ground truth value for each subject. Error bars illustrate the interquartile range (IQR) computed from 100 simulated data sets.

### Navajas et al. (2017), experiment 1

Figure 5a-d shows an analysis of data from subjects who performed either a perceptual or cognitive 2-AFC categorization task and additionally reported their confidence using a six-point rating scale. Data were collected by Navajas et al. (2017). Supplementary Figure 4 illustrates the model fit for an example subject by plotting the data in the format used in the original publication ^24^. Proportion correct and confidence level are plotted against stimulus variance. The confidence reports are split out by decision accuracy. The experiment consisted of 400 trials. To model these data, we used one lapse rate parameter (0%), one stimulus variance-specific sensitivity parameter (0.67, 0.53, 0.33, and 0.19), one decision criterion parameter (–0.59 degrees), one meta-uncertainty parameter (0.10), and five confidence criterion parameters (0.35, 1.19, 1.64, 2.29, and 3.36). The log-probability of the data under the model was –733.50.

**Supplementary Figure 4.**
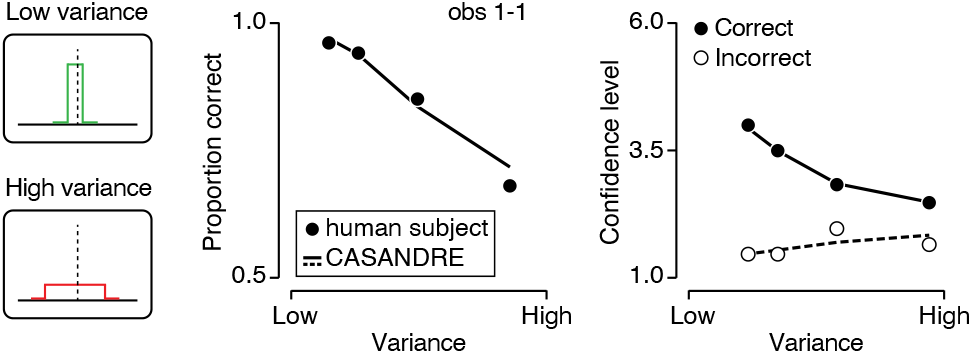
Model fit for an example subject from Navajas et al. (2017) (observer 1 in experiment 1). The subject judged whether the mean orientation of a sequence of 30 rapidly presented Gabor stimuli was tilted right or left. Left: Stimulus sequences were sampled from distributions with different orientation variance. Middle: Proportion correct choices is plotted against stimulus variance for an example subject. Right: Mean confidence level is plotted against stimulus variance, split out by decision accuracy. Symbols summarize observed choice behavior, the full line shows the fit of a two-stage process model of decision-making.

### Goodness-of-fit across datasets

Supplementary Figure 5 shows a comparison of the model predicted and observed choice behavior and confidence reports for the six tasks included in figure 7c.

**Supplementary Figure 5.**
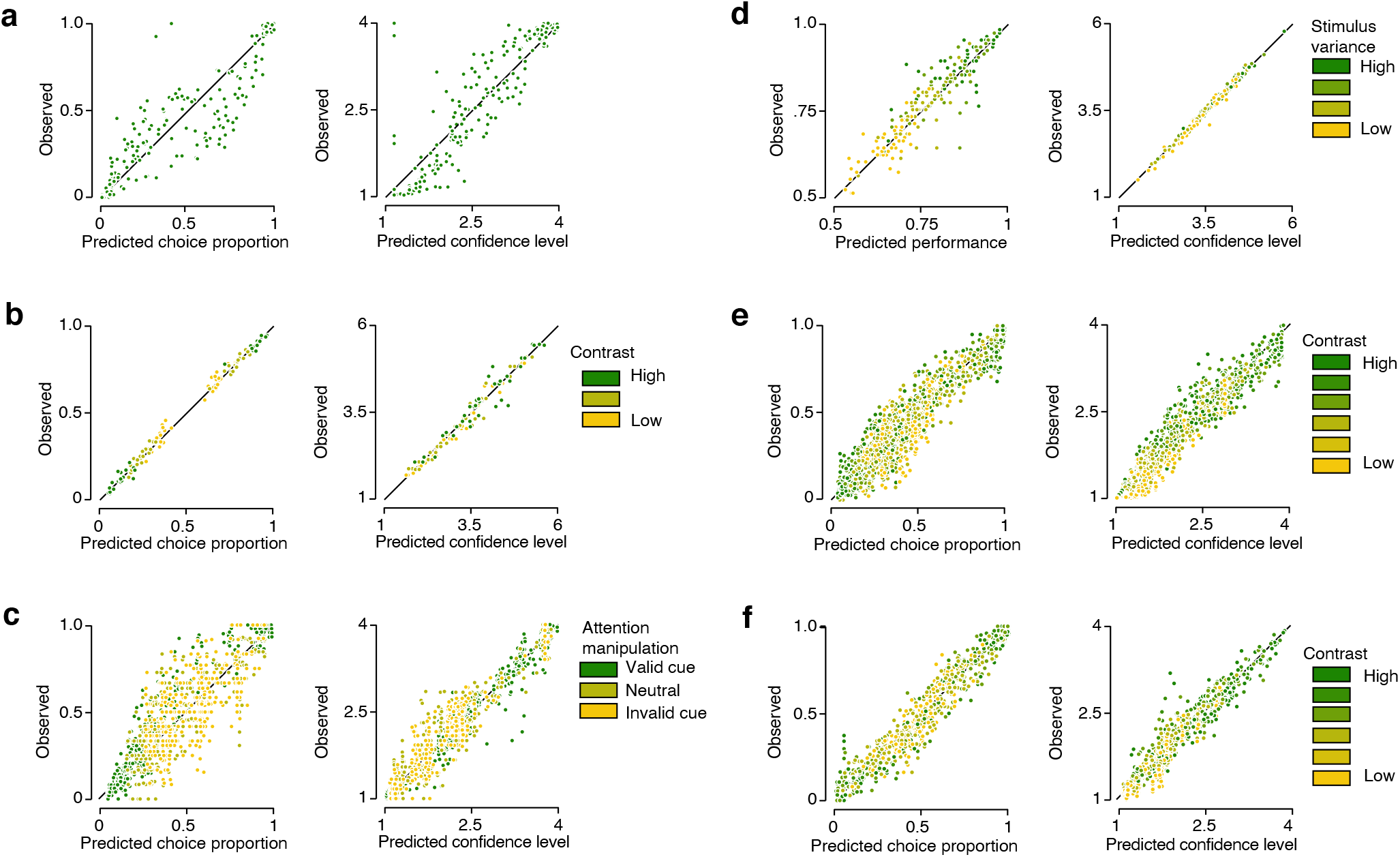
Each symbol summarizes a single stimulus condition for a single subject. Color indicates stimulus reliability. Left: Observed versus predicted choice behavior. Right: Observed versus predicted confidence level. (**a**) Data from Rausch et al. (2020): 25 subjects. (**b**) Shekhar and Rahnev (2021): 20 subjects. **c**) Denison et al. (2018): 12 subjects. **d**) Navajas et al. (2017): 50 subjects; **e**) Adler and Ma (2018) task 2: 34 subjects; **f**) Adler and Ma (2018) task 1: 19 subjects;

### Adler and Ma (2018), task 2

Figure 7c includes a data-point for task 2 from Adler and Ma (2018) and one for Denison et al. (2018). Both studies employed a task design in which subjects discriminated two categories of orientation distributions with the same mean but different standard deviations. Supplementary Figure 6 illustrates an example model fit for this task by plotting the psychometric function (top row) and accompanying confidence function (bottom row) for each stimulus contrast (columns). The subject completed 3,240 trials. To model these data, we used one lapse rate parameter (2.17%), one contrast-specific sensitivity parameter (0.68, 0.55, 0.46, 0.28, 0.20, and 0.07), one contrast-specific low decision criterion parameter (−4.05, -4.27, -4.35, -5.21, -7.30, and -11.34 degrees), one contrast-specific high decision criterion parameter (3.62, 3.76, 4.26, 5.36, 5.38, and 9.66 degrees), one meta-uncertainty parameter (0.69), three confidence criterion parameters for “Category A” choices (0.69, 1.15, and 1.89), and three confidence criterion parameters for “Category B” choices (0.65, 1.78, and 4.53). The log-probability of the data under the model was –4,842.5. (Note: while the model fit illustrated here employs asymmetric confidence criteria, all fits in Figure 7c were with symmetric confidence criteria for consistency.)

**Supplementary Figure 6.**
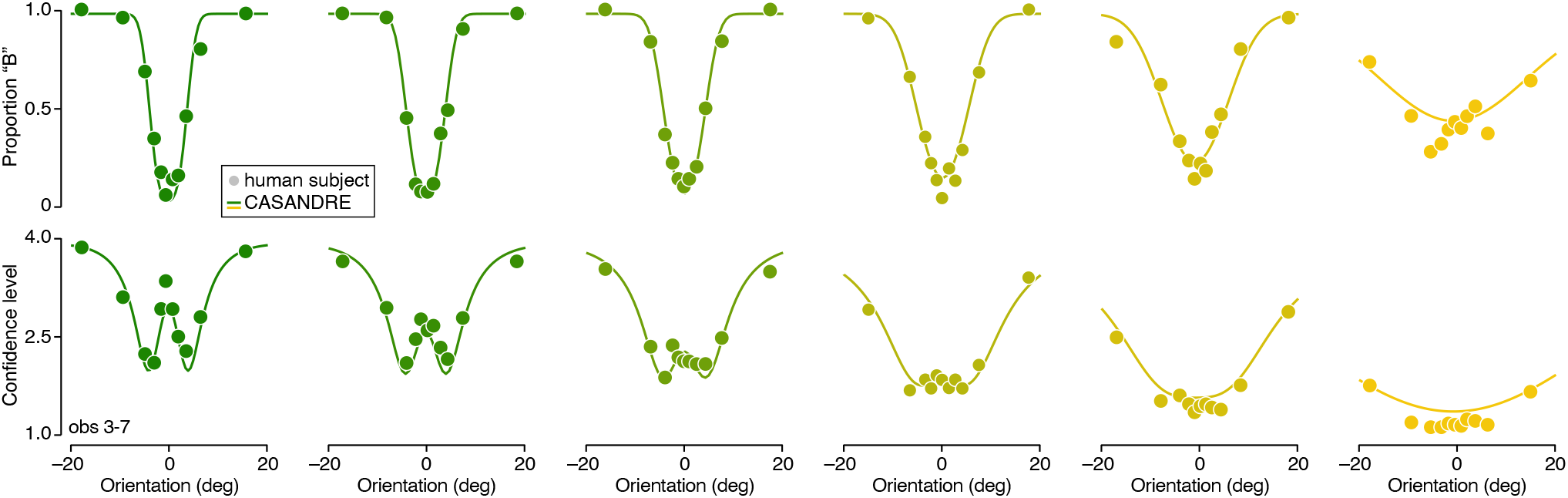
Model fits for an example subject from Adler and Ma (2018) (observer 7 in experiment 3). The subject judged whether a stimulus belonged to category A or B. Category A stimuli were drawn from a distribution with small orientation spread, category B stimuli were drawn from a distribution with large orientation spread. Stimuli varied in orientation and contrast. Top: Proportion of “Category B” choices is plotted against stimulus orientation, split out by stimulus contrast (columns, contrast decreases from left to right), for one example subject. Bottom: Same for mean confidence level. Symbols summarize observed choice behavior, the full lines show the fit of the CASANDRE model.

### Parameter trade-offs

Figure 4 illustrates a recovery analysis for the meta-uncertainty parameter of the CASANDRE model. Supplementary Figure 7 illustrates an additional analysis of the trade-off between meta-uncertainty and the other parameters of the CASANDRE model using the same generated data and model fits as in Figure 4c. Although the variance in meta-uncertainty explained by tradeoffs with confidence criterion can reach high levels, this is somewhat mitigated by denser stimulus sampling (Supplementary Fig. 7b, bottom) and is reasonable for datasets with a larger number of trials (Supplementary Fig. 7b, right) and for values of meta-uncertainty that are empirically observed more often (less than 1).

**Supplementary Figure 7.**
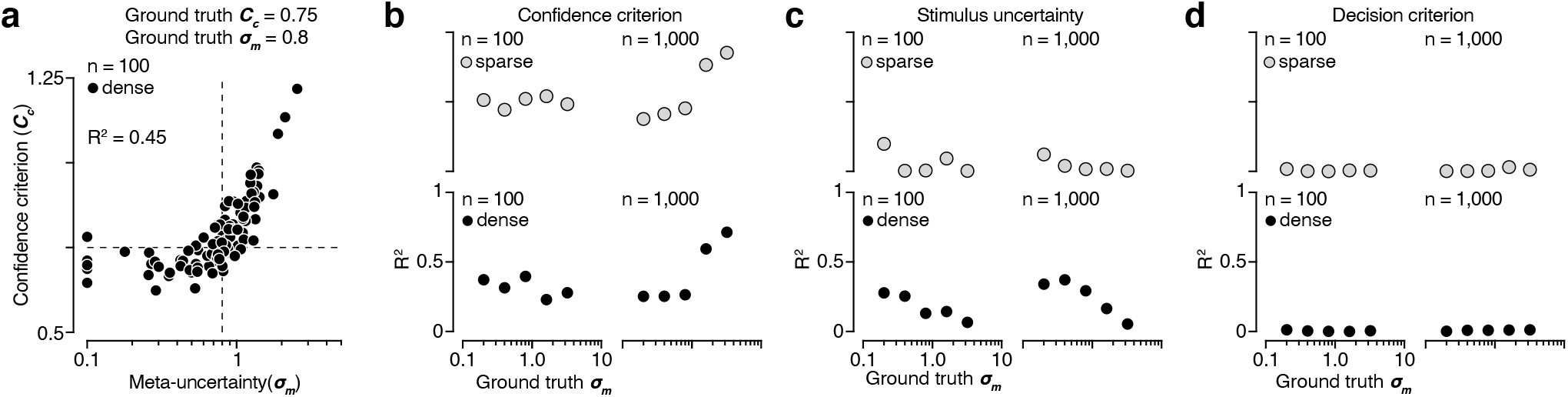
Trade-off between meta-uncertainty and other CASANDRE model parameters. (**a**) Parameter correlation for an example condition. Recovered meta-uncertainty and confidence criterion are plotted against each other for 100 model-generated datasets. The dashed lines represent the ground truth values for confidence criterion (*C*_*c*_ = 0.75) and meta-uncertainty (*σ*_*m*_ = 0.8). Each symbol represents one dataset generated with 100 trials and a dense stimulus sampling regime. (**b**) Trade-off between meta-uncertainty and confidence criterion. (**c**) Trade-off between meta-uncertainty and stimulus uncertainty. (**d** Trade-off between meta-uncertainty and decision criterion.)

### Comparison with a criteria-noise model

Shekhar and Rahnev recently described a hierarchical process model of confidence with desirable properties that dissociate a parameter capturing metacognitive ability from stimulus sensitivity and confidence reporting strategy ^28^. They refer to the parameter capturing metacognitive ability as “meta-noise” and find that log-normally distributed meta-noise provides a better quantitative and qualitative match to empirical data than normally distributed noise. In the CASANDRE model, the standard deviation of a log-normal distribution also serves as a metric for the metacognitive ability of an observer, however these two uses of log-normal noise, like the models themselves, are not equivalent. In the CASANDRE model, the confidence variable is distributed according to the ratio of a normally and log-normally distributed variable, whereas in the model of Shekhar and Rahnev the confidence variable has a normal distribution identical to the decision variable but the positions of the confidence criteria are subject to log-normally distributed noise. We thus refer to the model of Shekhar and Rahnev as the “criteria-noise” model.

We quantitatively compared the criteria-noise model with the CASANDRE model. Because the criteria-noise model is currently limited to experiments with two stimulus strengths, we did not apply it to data from ref. ^25^ (as we did for comparing other model variants in Fig. 3), but instead fit the CASANDRE model to the data reported in their original paper ^28^. For purposes of quantitative comparison to the criteria-noise model, we fit the CASANDRE model with asymmetric confidence criteria (yielding 15 total parameters, see Methods). First, we compared the CASANDRE model to the criteria-noise model as described by Shekhar and Rhanev, with a different set of confidence criteria for each of three contrast values (yielding 35 total parameters). The CASANDRE model significantly outperformed the criteria-noise model (median difference in AIC = 7.8; *P* = 0.002, Wilcoxon signed-rank test; Supplementary Fig. 8a, top). Second, we compared the CASANDRE model to a simpler version of the criteria-noise model with only one set of confidence critiera (yielding 15 total parameters). There was no difference in performance between the CASANDRE model and this variant of the criteria-noise model (median difference in AIC = -0.1; *P* = 1, Wilcoxon signed-rank test; Supplementary Fig. 8a, bottom). Note: we discovered an incorrect scaling of likelihood values in the original code accompanying ref. ^28^. We fixed this scaling and thus the AIC values used in this model comparison for the criteria-noise model differ from those reported in ref. ^28^.

Shekhar and Rahnev demonstrated that the level of confidence criteria noise can serve as a measure of metacognitive ability uncontaminated by stimulus sensitivity or confidence reporting strategy ^28^. For comparison, we performed the same analysis using the CASANDRE model. Following their procedure, we mapped each subject’s continuous confidence reports into five different binary confidence rating scales, biasing confident reports to be more liberal or conservative. For each subject, we fit the CASANDRE model independently to each of these five remapped datasets across the three contrast levels. We removed one subject that in some conditions did not generate a single response to one of the four possible response options. Meta-uncertainty was largely insensitive to both confidence reporting strategy and stimulus sensitivity (Supplementary Fig. 8b; compare to Fig. 11a in Shekhar and Rahnev ^28^). A two-way ANOVA revealed no main effect of confidence criterion (F(4, 18) = 1.97, *P* = 0.11) or stimulus contrast (F(2, 18) = 0.04, *P* = 0.96) on meta-uncertainty. Further, the interaction between confidence reporting strategy and stimulus sensitivity was not significant (F(2, 4) = 1.57, *P* = 0.14). These results along with the model comparison (Supplementary Fig. 8a) demonstrate that the CASANDRE model performs quantitatively at least as well as the criteria-noise model in explaining the data reported in ref. ^28^.

We now turn to several more qualitative considerations that favor the CASANDRE model compared with the criteria-noise model. First, Shekhar and Rahnev show that empirical, averaged zROC functions have significant curvature compared with the straight zROC functions predicted by signal detection theory (their Fig. 4 and 5b). The criteria-noise model shows curved zROC functions but, as the authors note, they resemble piecewise linear functions rather than the smoothly curving zROC functions of the empirical data (see their Fig. 11, bottom left). The CASANDRE model generates smoothly curving averaged zROC functions that more closely resemble the empirically estimated zROC curves (Supplementary Fig. 8c; compare with Fig. 4 and 5b in ref. ^28^). Second, the CASANDRE model is easier to fit to data given that its parameters can be optimized using standard maximum likelihood estimation procedures, rather than the purpose-built, two-stage parameter search algorithm developed by Shekhar and Rahnev. Third, the CASANDRE model is more general and can be applied to experiments that vary stimulus strength in addition to stimulus uncertainty (such as ref. ^25^), whereas the criteria-noise model is limited to experiments with two stimulus strengths. This is because the criteria-noise model makes the assumption that confidence is measured in units of *d*′, but does not specify the computation that transforms units of stimulus to units of *d*′. Fourth, the CASANDRE model specifies this confidence computation and posits that it is exactly noise in this transformation that can lead to limited metacognitive ability. Analogous to stimulus discrimination ability being limited by variation in the estimation of the stimulus, the CASANDRE model posits that metacognition is limited by variation in the estimation of the uncertainty required to compute confidence. In contrast, the criteria-noise model posits that lower metacognitive ability arises from the inability of subjects to maintain constant confidence criteria. Stochastic confidence criteria cause problems for model tractability, allowing for them to cross both the decision criterion and each other. By casting criteria-noise as log-normally distributed, Shekhar and Rahnev avoid the problem of crossovers with the decision criterion, but to solve the problem of crossovers between confidence criteria they make the questionable assumption that noise is perfectly correlated across criteria. The CASANDRE model naturally avoids both of these issues. Fifth, the process captured by the CASANDRE model leads to new predictions about how metacognitive ability can be experimentally manipulated. Figure 7 illustrates how increasing the number of uncertainty levels in a task increases meta-uncertainty. If an inability to maintain stable criteria were a source of lower metacognitive ability instead, increasing the number of confidence response levels on the rating scale used by subjects should lower metacognitive ability (and increase meta-uncertainty estimated from the CASANDRE model). We see no evidence for this prediction when rearranging the estimated meta-uncertainty across tasks according to the number confidence response levels(Supplementary Fig. 8d).

**Supplementary Figure 8.**
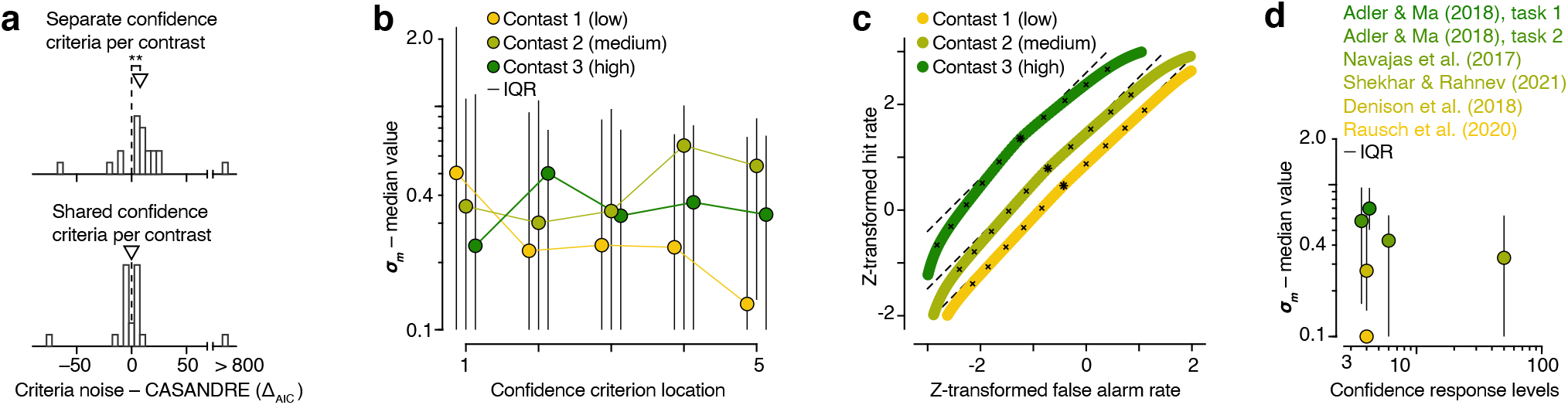
Comparison with Shekhar and Rahnev (2021). (**a**) Distribution of the difference in AIC value across 20 subjects for the CASANDRE model compared with the criteria-noise model with a different set of confidence criteria for each of three contrast levels (top) or with one shared set of confidence criteria across all three contrasts (bottom). Positive values indicate evidence favoring the CASANDRE model. Arrows indicate the median of the distribution. ** P < 0.01, Wilcoxon signed-rank test. (**b**) Median meta-uncertainty across 20 subjects estimated independently for each contrast and confidence criterion location. Error bars illustrate the interquartile range (IQR) across subjects. Compare to Fig. 11a in ref. ^28^. (**c**) Averaged zROC functions across 20 subjects generated from fits of the CASANDRE model. The location of the decision criterion is indicated by an asterisk, and the location of each confidence criterion is indicated by an x. The dashed lines illustrate the linear zROC functions predicted from signal detection theory. Compare to Fig. 4, 5b, and 10 in ref. ^28^. (**d**) Median level of meta-uncertainty plotted against number of confidence response levels for six confidence experiments. Error bars illustrate the interquartile range (IQR) across subjects. Note the symbol representing ref. ^28^ is plotted at 50 although subjects rated their confidence on a continuous scale ranging from 50-100.

## References

1. Florent Meyniel, Mariano Sigman, and Zachary F. Mainen. Confidence as Bayesian Probability: From Neural Origins to Behavior. Neuron, 88(1):78–92, October 2015.

2. Jan Drugowitsch, André G. Mendonça, Zachary F. Mainen, and Alexandre Pouget. Learning optimal decisions with confidence. Proceedings of the National Academy of Sciences, 116(49):24872–24880, December 2019.

3. Braden A. Purcell and Roozbeh Kiani. Hierarchical decision processes that operate over distinct timescales underlie choice and changes in strategy. Proceedings of the National Academy of Sciences, 113(31):E4531–E4540, August 2016.

4. Bahador Bahrami, Karsten Olsen, Peter E. Latham, Andreas Roepstorff, Geraint Rees, and Chris D. Frith. Optimally Interacting Minds. Science, 329(5995):1081–1085, August 2010.

5. Charles Sanders Peirce and Joseph Jastrow. On small differences in sensation. Memoirs of the National Academy of Sciences, 3, 1884.

6. Roger Ratcliff. A theory of memory retrieval. Psychological Review, 85(2):59–108, 1978.

7. Douglas Vickers. Decision processes in visual perception. Academic Press, New York, 1979.

8. Vincent de Gardelle, François Le Corre, and Pascal Mamassian. Confidence as a Common Currency between Vision and Audition. PLOS ONE, 11(1):e0147901, January 2016.

9. Stephen M. Fleming, Rimona S. Weil, Zoltan Nagy, Raymond J. Dolan, and Geraint Rees. Relating Introspective Accuracy to Individual Differences in Brain Structure. Science, 329(5998):1541–1543, September 2010.

10. Marion Rouault, Tricia Seow, Claire M. Gillan, and Stephen M. Fleming. Psychiatric Symptom Dimensions Are Associated With Dissociable Shifts in Metacognition but Not Task Performance. Biological Psychiatry, 84(6):443–451, September 2018.

11. Deanna Kuhn. Theory of mind, metacognition, and reasoning: A life-span perspective. In Children’s reasoning and the mind, pages 301–326. Psychology Press, 2000.

12. Thomas O. Nelson. A comparison of current measures of the accuracy of feeling-of-knowing predictions. Psychological Bulletin, 95(1):109–133, 1984.

13. Stephen M. Fleming and Hakwan C. Lau. How to measure metacognition. Frontiers in Human Neuroscience, 8:443, 2014.

14. Pascal Mamassian. Visual Confidence. Annual Review of Vision Science, 2(1):459–481, 2016.

15. Matthias Guggenmos. Measuring metacognitive performance: type 1 performance dependence and test-retest reliability. Neuroscience of Consciousness, 2021(1):iab040, December 2021.

16. L. Festinger. Studies in decision: I. Decision-time, relative frequency of judgment and subjective confidence as related to physical stimulus difference. Journal of Experimental Psychology, 32(4):291–306, 1943.

17. Jinoos Hosseini and William R. Ferrell. Detectability of correctness: A measure of knowing that one knows. Instructional Science, 11(2):113–127, August 1982.

18. Thomas S. Critchfield. Signal-Detection Properties of Verbal Self-Reports. Journal of the Experimental Analysis of Behavior, 60(3):495–514, 1993.

19. Susan J. Galvin, John V. Podd, Vit Drga, and John Whitmore. Type 2 tasks in the theory of signal detectability: Discrimination between correct and incorrect decisions. Psychonomic Bulletin & Review, 10(4):843–876, December 2003.

20. Baptiste Caziot and Pascal Mamassian. Perceptual confidence judgments reflect self-consistency. Journal of Vision, 21(12):8, November 2021.

21. A. Pouget, J. Drugowitsch, and A. Kepecs. Confidence and certainty: distinct probabilistic quantities for different goals. Nature neuroscience, 19(3):366–374, 2016.

22. David Marvin Green and John A. Swets. Signal detection theory and psychophysics, volume 1. Wiley New York, 1966.

23. Dobromir Rahnev, Kobe Desender, Alan L. F. Lee, William T. Adler, David Aguilar-Lleyda, Başak Akdoğ an, Polina Arbuzova, Lauren Y. Atlas, Fuat Balci, Ji Won Bang, Indrit Bègue, Damian P. Birney, Timothy F. Brady, Joshua Calder-Travis, Andrey Chetverikov, Torin K. Clark, Karen Davranche, Rachel N. Denison, Troy C. Dildine, Kit S. Double, Yalçin A. Duyan, Nathan Faivre, Kaitlyn Fallow, Elisa Filevich, Thibault Gajdos, Regan M. Gallagher, Vincent de Gardelle, Sabina Gherman, Nadia Haddara, Marine Hainguerlot, Tzu-Yu Hsu, Xiao Hu, Iñaki Iturrate, Matt Jaquiery, Justin Kantner, Marcin Koculak, Mahiko Konishi, Christina Koß, Peter D. Kvam, Sze Chai Kwok, Maël Lebreton, Karolina M. Lempert, Chien Ming Lo, Liang Luo, Brian Maniscalco, Antonio Martin, Sébastien Massoni, Julian Matthews, Audrey Mazancieux, Daniel M. Merfeld, Denis O’Hora, Eleanor R. Palser, Borysław Paulewicz, Michael Pereira, Caroline Peters, Marios G. Philiastides, Gerit Pfuhl, Fernanda Prieto, Manuel Rausch, Samuel Recht, Gabriel Reyes, Marion Rouault, Jérôme Sackur, Saeedeh Sadeghi, Jason Samaha, Tricia X. F. Seow, Medha Shekhar, Maxine T. Sherman, Marta Siedlecka, Zuzanna Skóra, Chen Song, David Soto, Sai Sun, Jeroen J. A. van Boxtel, Shuo Wang, Christoph T. Weidemann, Gabriel Weindel, Michał Wierzchoń, Xinming Xu, Qun Ye, Jiwon Yeon, Futing Zou, and Ariel Zylberberg. The Confidence Database. Nature Human Behaviour, 4(3):317–325, March 2020.

24. Joaquin Navajas, Chandni Hindocha, Hebah Foda, Mehdi Keramati, Peter E. Latham, and Bahador Bahrami. The idiosyncratic nature of confidence. Nature Human Behaviour, 1(11):810–818, November 2017.

25. William T. Adler and Wei Ji Ma. Comparing Bayesian and non-Bayesian accounts of human confidence reports. PLOS Computational Biology, 14(11):e1006572, November 2018.

26. Rachel N. Denison, William T. Adler, Marisa Carrasco, and Wei Ji Ma. Humans incorporate attention-dependent uncertainty into perceptual decisions and confidence. Proceedings of the National Academy of Sciences, 115(43):11090–11095, October 2018.

27. Manuel Rausch, Michael Zehetleitner, Marco Steinhauser, and Martin E. Maier. Cognitive modelling reveals distinct electrophysiological markers of decision confidence and error monitoring. NeuroImage, 218:116963, September 2020.

28. Medha Shekhar and Dobromir Rahnev. The nature of metacognitive inefficiency in perceptual decision making. Psychological Review, 128(1):45–70, 2021.

29. J. D. Balakrishnan and Roger Ratcliff. Testing models of decision making using confidence ratings in classification. Journal of Experimental Psychology: Human Perception and Performance, 22(3):615–633, 1996.

30. William R. Ferrell. A model for realism of confidence judgments: Implications for underconfidence in sensory discrimination. Perception & Psychophysics, 57(2):246–254, January 1995.

31. Adam Kepecs, Naoshige Uchida, Hatim A. Zariwala, and Zachary F. Mainen. Neural correlates, computation and behavioural impact of decision confidence. Nature, 455(7210):227–231, September 2008.

32. Michel Treisman and Andrew Faulkner. The setting and maintenance of criteria representing levels of confidence. Journal of Experimental Psychology: Human Perception and Performance, 10(1):119–139, 1984.

33. Thomas S. Wallsten and Claudia González-Vallejo. Statement verification: A stochastic model of judgment and response. Psychological Review, 101(3):490–504, 1994.

34. E. T. Jaynes. Information Theory and Statistical Mechanics. Physical Review, 106(4):620–630, May 1957.

35. Sanjoy Mahajan. Street-Fighting Mathematics: The Art of Educated Guessing and Opportunistic Problem Solving. The MIT Press, 2010.

36. Shannon M. Locke, Elon Gaffin-Cahn, Nadia Hosseinizaveh, Pascal Mamassian, and Michael S. Landy. Priors and payoffs in confidence judgments. Attention, Perception, & Psychophysics, 82(6):3158–3175, August 2020.

37. Andra Mihali, Marianne Broeker, and Guillermo Horga. Insightful inference compensates for distorted perception. bioRxiv, page 2021.11.13.468497, November 2021. Type: article.

38. Christopher R. Fetsch, Roozbeh Kiani, William T. Newsome, and Michael N. Shadlen. Effects of cortical microstimulation on confidence in a perceptual decision. Neuron, 83(4):797–804, 2014.

39. Christopher R Fetsch, Naomi N Odean, Danique Jeurissen, Yasmine El-Shamayleh, Gregory D Horwitz, and Michael N Shadlen. Focal optogenetic suppression in macaque area MT biases direction discrimination and decision confidence, but only transiently. eLife, 7:e36523, July 2018.

40. Kai Xue, Medha Shekhar, and Dobromir Rahnev. Examining the robustness of the relationship between metacognitive efficiency and metacognitive bias. Consciousness and Cognition, 95:103196, October 2021.

41. Ji Won Bang, Medha Shekhar, and Dobromir Rahnev. Sensory noise increases metacognitive efficiency. Journal of Experimental Psychology. General, 148(3):437–452, March 2019.

42. Li Yan McCurdy, Brian Maniscalco, Janet Metcalfe, Ka Yuet Liu, Floris P. de Lange, and Hakwan Lau. Anatomical Coupling between Distinct Metacognitive Systems for Memory and Visual Perception. Journal of Neuroscience, 33(5):1897–1906, January 2013.

43. Benjamin Baird, Matthew Cieslak, Jonathan Smallwood, Scott T. Grafton, and Jonathan W. Schooler. Regional White Matter Variation Associated with Domain-specific Metacognitive Accuracy. Journal of Cognitive Neuroscience, 27(3):440–452, March 2015.

44. Alan L. F. Lee, Eugene Ruby, Nathan Giles, and Hakwan Lau. Cross-Domain Association in Metacognitive Efficiency Depends on First-Order Task Types. Frontiers in Psychology, 9, 2018.

45. Brian Maniscalco and Hakwan Lau. A signal detection theoretic approach for estimating metacognitive sensitivity from confidence ratings. Consciousness and Cognition, 21(1):422–430, March 2012.

46. William Kruskal. Relative Importance by Averaging over Orderings. The American Statistician, 41(1):6–10, February 1987.

47. Ulrike Grömping. Estimators of Relative Importance in Linear Regression Based on Variance Decomposition. The American Statistician, 61(2):139–147, May 2007.

48. Wendy E. Shields, J. David Smith, Katarina Guttmannova, and David A. Washburn. Confidence Judgments by Humans and Rhesus Monkeys. The Journal of general psychology, 132(2):165–186, April 2005.

49. Shannon M. Locke, Michael S. Landy, and Pascal Mamassian. Suprathreshold perceptual decisions constrain models of confidence. Technical report, PsyArXiv, December 2021. type: article.

50. Dobromir Rahnev, Tarryn Balsdon, Lucie Charles, Vincent de Gardelle, Rachel N. Denison, Kobe Desender, Nathan Faivre, Elisa Filevich, Stephen Fleming, Janneke Jehee, Hakwan Lau, Alan L. F. Lee, Shannon M. Locke, Pascal Mamassian, Brian Odegaard, Megan A. K. Peters, Gabriel Reyes, Marion Rouault, Jérôme Sackur, Jason Samaha, Claire Sergent, Maxine Sherman, Marta Siedlecka, David Soto, Alexandra Vlassova, and Ariel Zylberberg. Consensus goals for the field of visual metacognition. Technical report, PsyArXiv, April 2021. type: article.

51. Yoshiaki Ko and Hakwan Lau. A detection theoretic explanation of blindsight suggests a link between conscious perception and metacognition. Philosophical Transactions of the Royal Society B: Biological Sciences, 367(1594):1401–1411, May 2012.

52. Yutaka Komura, Akihiko Nikkuni, Noriko Hirashima, Teppei Uetake, and Aki Miyamoto. Responses of pulvinar neurons reflect a subject’s confidence in visual categorization. Nature Neuroscience, 16(6):749–755, June 2013.

53. Sébastien Massoni, Thibault Gajdos, and Jean-Christophe Vergnaud. Confidence measurement in the light of signal detection theory. Frontiers in Psychology, 5, 2014.

54. Ariel Zylberberg, Pablo Barttfeld, and Mariano Sigman. The construction of confidence in a perceptual decision. Frontiers in Integrative Neuroscience, 6, 2012.

55. Brian Maniscalco, Megan A. K. Peters, and Hakwan Lau. Heuristic use of perceptual evidence leads to dissociation between performance and metacognitive sensitivity. Attention, Perception, & Psychophysics, 78(3):923–937, April 2016.

56. Megan A. K. Peters, Thomas Thesen, Yoshiaki D. Ko, Brian Maniscalco, Chad Carlson, Matt Davidson, Werner Doyle, Ruben Kuzniecky, Orrin Devinsky, Eric Halgren, and Hakwan Lau. Perceptual confidence neglects decision-incongruent evidence in the brain. Nature Human Behaviour, 1(7):1–8, July 2017.

57. Stephen M. Fleming and Nathaniel D. Daw. Self-evaluation of decision-making: A general Bayesian framework for metacognitive computation. Psychological Review, 124(1):91–114, 2017.

58. Christopher R. Fetsch, Roozbeh Kiani, and Michael N. Shadlen. Predicting the accuracy of a decision: a neural mechanism of confidence. In Cold Spring Harbor symposia on quantitative biology, volume 79, pages 185–197. Cold Spring Harbor Laboratory Press, 2014.

59. Peter R Murphy, Ian H Robertson, Siobhán Harty, and Redmond G O’Connell. Neural evidence accumulation persists after choice to inform metacognitive judgments. eLife, 4:e11946, December 2015.

60. Koosha Khalvati, Roozbeh Kiani, and Rajesh P. N. Rao. Bayesian inference with incomplete knowledge explains perceptual confidence and its deviations from accuracy. Nature Communications, 12(1):5704, September 2021.

61. Brian Maniscalco and Hakwan Lau. The signal processing architecture underlying subjective reports of sensory awareness. Neuroscience of Consciousness, 2016(1), January 2016.

62. Armin Lak, Gil M. Costa, Erin Romberg, Alexei A. Koulakov, Zachary F. Mainen, and Adam Kepecs. Orbitofrontal Cortex Is Required for Optimal Waiting Based on Decision Confidence. Neuron, 84(1):190–201, October 2014.

63. Roozbeh Kiani and Michael N. Shadlen. Representation of confidence associated with a decision by neurons in the parietal cortex. science, 324(5928):759–764, 2009.

64. Joshua I. Sanders, Balázs Hangya, and Adam Kepecs. Signatures of a Statistical Computation in the Human Sense of Confidence. Neuron, 90(3):499–506, May 2016.

65. Balázs Hangya, Joshua I. Sanders, and Adam Kepecs. A Mathematical Framework for Statistical Decision Confidence. Neural Computation, 28(9):1840–1858, September 2016.

66. William T. Adler and Wei Ji Ma. Limitations of Proposed Signatures of Bayesian Confidence. Neural Computation, 30(12):3327–3354, December 2018.

67. Hsin-Hung Li and Wei Ji Ma. Confidence reports in decision-making with multiple alternatives violate the Bayesian confidence hypothesis. Nature Communications, 11(1):2004, April 2020.

68. Laura S. Geurts, James R. H. Cooke, Ruben S. van Bergen, and Janneke F. M. Jehee. Subjective confidence reflects representation of Bayesian probability in cortex. Nature Human Behaviour, pages 1–12, January 2022.

69. Benedetto De Martino, Stephen M. Fleming, Neil Garrett, and Raymond Dolan. Confidence in value-based choice. Nature neuroscience, 16(1):105–110, January 2013.

70. Marc O. Ernst and Martin S. Banks. Humans integrate visual and haptic information in a statistically optimal fashion. Nature, 415(6870):429, 2002.

71. Christopher R. Fetsch, Alexandre Pouget, Gregory C. DeAngelis, and Dora E. Angelaki. Neural correlates of reliability-based cue weighting during multisensory integration. Nature neuroscience, 15(1):146, 2012.

72. Ahmad T. Qamar, R. James Cotton, Ryan G. George, Jeffrey M. Beck, Eugenia Prezhdo, Allison Laudano, Andreas S. Tolias, and Wei Ji Ma. Trial-to-trial, uncertainty-based adjustment of decision boundaries in visual categorization. Proceedings of the National Academy of Sciences, 110(50):20332–20337, December 2013.

73. Wei Ji Ma, Jeffrey M. Beck, Peter E. Latham, and Alexandre Pouget. Bayesian inference with probabilistic population codes. Nature neuroscience, 9(11):1432, 2006.

74. Gergő Orbán, Pietro Berkes, József Fiser, and Máté Lengyel. Neural variability and sampling-based probabilistic representations in the visual cortex. Neuron, 92(2):530–543, 2016.

75. Ruben S. van Bergen, Wei Ji Ma, Michael S. Pratte, and Janneke F. M. Jehee. Sensory uncertainty decoded from visual cortex predicts behavior. Nature Neuroscience, 18(12):1728–1730, December 2015.

76. Olivier J. Hénaff, Zoe M. Boundy-Singer, Kristof Meding, Corey M. Ziemba, and Robbe L. T. Goris. Representation of visual uncertainty through neural gain variability. Nature Communications, 11(1):2513, May 2020.

77. Edgar Y. Walker, R. James Cotton, Wei Ji Ma, and Andreas S. Tolias. A neural basis of probabilistic computation in visual cortex. Nature Neuroscience, 23(1):122–129, January 2020.

78. Dylan Festa, Amir Aschner, Aida Davila, Adam Kohn, and Ruben Coen-Cagli. Neuronal variability reflects probabilistic inference tuned to natural image statistics. Nature Communications, 12(1):3635, June 2021.

79. Ariel Zylberberg, Pieter R. Roelfsema, and Mariano Sigman. Variance misperception explains illusions of confidence in simple perceptual decisions. Consciousness and Cognition, 27:246–253, July 2014.

80. Ariel Zylberberg, Christopher R Fetsch, and Michael N Shadlen. The influence of evidence volatility on choice, reaction time and confidence in a perceptual decision. eLife, 5:e17688, October 2016.

81. Santiago Herce Castañón, Rani Moran, Jacqueline Ding, Tobias Egner, Dan Bang, and Christopher Summerfield. Human noise blindness drives suboptimal cognitive inference. Nature Communications, 10(1):1719, April 2019.

82. John Palmer, Alexander C. Huk, and Michael N. Shadlen. The effect of stimulus strength on the speed and accuracy of a perceptual decision. Journal of Vision, 5(5):1, May 2005.

83. Timothy D. Hanks, Mark E. Mazurek, Roozbeh Kiani, Elisabeth Hopp, and Michael N. Shadlen. Elapsed Decision Time Affects the Weighting of Prior Probability in a Perceptual Decision Task. Journal of Neuroscience, 31(17):6339–6352, April 2011.

84. Roozbeh Kiani, Leah Corthell, and Michael N. Shadlen. Choice Certainty Is Informed by Both Evidence and Decision Time. Neuron, 84(6):1329–1342, December 2014.

85. John Gibbon. Scalar expectancy theory and Weber’s law in animal timing. Psychological Review, 84(3):279–325, 1977.

86. Mehrdad Jazayeri and Michael N. Shadlen. Temporal context calibrates interval timing. Nature Neuroscience, 13(8):1020– 1026, August 2010.

87. Mehrdad Jazayeri and Michael N. Shadlen. A Neural Mechanism for Sensing and Reproducing a Time Interval. Current Biology, 25(20):2599–2609, October 2015.

88. Wilson S. Geisler. Ideal observer analysis. In The Visual Neurosciences, volume 10, pages 825–837. MIT Press, Boston, 2003.

89. Yair Weiss, Eero P. Simoncelli, and Edward H. Adelson. Motion illusions as optimal percepts. Nature neuroscience, 5(6):598, 2002.

90. Navindra Persaud, Peter McLeod, and Alan Cowey. Post-decision wagering objectively measures awareness. Nature Neuroscience, 10(2):257–261, February 2007.

91. Zoltán Dienes and Anil Seth. Gambling on the unconscious: A comparison of wagering and confidence ratings as measures of awareness in an artificial grammar task. Consciousness and Cognition, 19(2):674–681, June 2010.

92. Zahra Murad, Martin Sefton, and Chris Starmer. How do risk attitudes affect measured confidence? Journal of Risk and Uncertainty, 52(1):21–46, February 2016.

93. F. A. Wichmann and N. J. Hill. The psychometric function: I. Fitting, sampling, and goodness of fit. Perception & Psychophysics, 63(8):1293–1313, November 2001.

